# MucD regulates alginate biosynthesis through the proteolytic control of AlgX and AlgK in *Pseudomonas aeruginosa*

**DOI:** 10.64898/2026.08.29.748010

**Authors:** Yujun Jiang, Xin-Fu Yan, Rya Ero, Chao Wang, Kanaga Sabapathy, Yong-Gui Gao

**Author notes:** Correspondence (Y.-G.G.).

## Abstract

*Pseudomonas aeruginosa* is an opportunistic human pathogen capable of infecting a wide range of tissues and organs. Its persistence during chronic infection is strongly associated with biofilm formation, which depends on extracellular polysaccharides such as alginate. The HtrA-like periplasmic serine protease MucD is a key regulator of bacterial virulence, stress response, and alginate production, yet its molecular mechanism has remained largely unclear. Here, we discovered the alginate acetylation and export proteins AlgX and AlgK as MucD substrates, and characterized their degradation by mass spectrometry and bioinformatic analysis. We further determined the cryo-EM structure of MucD bound to an AlgK-derived substrate peptide, offering atomic insights into MucD oligomerization assembly, substrate recognition, and specificity. Together with structure-guided mutagenesis and biochemical assays, our results revealed that MucD proteolytic activity is governed by an equilibrium between a resting 12-mer and an active trimer. Crucially, we demonstrate that MucD represses alginate biosynthesis post-translationally, in addition to its previously implicated role in transcriptional regulation. These findings define a distinct activation mechanism and regulatory function for MucD and provide new insight into bacterial HtrA-like serine proteases.

## Introduction

*Pseudomonas aeruginosa* is a major opportunistic human pathogen and a frequent cause of nosocomial infections, particularly in immunocompromised patients [1]. Its pathogenicity is supported by the production of diverse virulence factors, including extracellular polysaccharides (EPS). EPS are key components of bacterial biofilm, which shields bacteria from antibiotics, oxidative stress, and host immune defenses during chronic infections [2]. Alginate, together with the EPS components Pel and Psl, plays an important role in biofilm formation and maturation. Alginate is synthesized through a synthase-dependent pathway encoded by 12-gene *alg* operon [3]. Transcription of the *alg* operon is tightly controlled by the sigma factor AlgU/T, and mutations affecting AlgU/T or its regulatory proteins are frequently found in clinical alginate-overproducing (mucoid) isolates [4].

Protease-mediated protein quality control is essential for cellular homeostasis, particularly under environmental stress [5]. The high-temperature requirement A (HtrA) family of serine proteases are conserved from bacteria to humans and contribute to stress adaptation by degrading misfolded or damaged proteins [6]. HtrA-like proteases typically contain a structurally conserved protease domain and one or two PDZ (**p**ost synaptic density protein 95 (PSD95), *Drosophila* **d**isc large tumor suppressor (Dlg1), and **z**onula occludens-1 protein (zo-1)) domains [7], which mediate protein-protein interactions and substrate binding [8]. In Gram-negative bacteria, HtrA-like proteases have attracted interest as potential antimicrobial targets because of their important physiological roles, conserved architecture, and defined substrate-binding sites [9]. For example, small-molecule inhibition of HtrA-like protease DegS suppresses stress-response activation and antibiotic resistance in *E. coli* [10]. DegP, another HtrA-like protease in Gram-negative bacteria, is central to periplasmic protein homeostasis [11]. Notably, its activity is tightly controlled by oligomerization: substrate binding converts an inactive hexameric resting state into larger, proteolytically active 12-mer and 24-mer assemblies [12].

The *mucD* gene is located within an autoregulated operon comprising *algU*, *mucA*, *mucB*, *mucC,* and *mucD* in *P. aeruginosa* [13]. MucD is an HtrA-like periplasmic protease and DegP homologue found in the genus *Pseudomonas* [14]. It contains a protease domain followed by two C-terminal PDZ domains (PDZ1 and PDZ2), and has been implicated in bacterial response to heat stress and reactive oxygen species [15,16]. Loss of *mucD* attenuates virulence in infection models of *Arabidopsis*, *C. elegans,* and mice [16,17]. MucD also negatively regulates alginate biosynthesis at the transcriptional level by indirectly repressing AlgU/T activity [18]. Affinity pull-down studies further suggested that MucD could form a periplasmic complex with the terminal acetyltransferase AlgX and tetratricopeptide repeat (TPR)-rich scaffold protein AlgK [14]. Both AlgK and AlgX are required for alginate chain maturation, modification, and secretion [19].

Despite the established roles of MucD in bacterial stress response, virulence, and alginate biosynthesis, its proteolytic function, physiological substrates, and structural basis of substrate recognition remain poorly understood. To date, the only reported MucD structure is that of a truncated *P. syringe* MucD lacking the PDZ2 domain [20]. Here we identified AlgK and AlgX as MucD substrates, determined the cryo-EM structure of substrate-bound full-length MucD, and characterized its proteolytic properties, oligomerization mode, substrate recognition and specificity, as well as physiological role in alginate biosynthesis. Together, our results define the functional and structural characteristics of MucD and offer atomic insights into HtrA-like protease-mediated proteolytic regulation in *P. aeruginosa*.

## Results

### Alginate biosynthesis pathway proteins AlgK and AlgX are substrates of MucD protease

Among alginate biosynthesis proteins, AlgK acts as a structural scaffold, tethering the inner-membrane alginate polymerase to the outer-membrane pore [21]; AlgX is an essential periplasmic acetyltransferase that O-acetylates the polymer, a modification required for alginate gel-like properties and resistance to host defenses [22]. Together, they form the AlgKX complex, which ensures proper alginate maturation and its transit through the periplasm prior to secretion [19]. A previous study reported a large periplasmic complex involving MucD, AlgK, and AlgX [14], but whether AlgK and AlgX are MucD substrates remained unknown. To address this question, MucD was expressed and purified from the wild-type *P. aeruginosa* PAO1 strain. The initial size-exclusion chromatography (SEC) profile showed that MucD primarily exists in two oligomeric states (Fig. 1A). Isolation and re-application of two oligomeric populations to analytical SEC separately further identified them as a trimer and a 12-mer (main population), based on molecular weight (MW) calibration (Fig. S1 A and B). Analytical SEC also indicated that the MucD 12-mer and trimer exist in a dynamic equilibrium in solution. Furthermore, no convincing additional protein co-eluting with MucD was detected based on SDS-PAGE (Fig. 1A) or MALDI-TOF/MS (Table S1). We next tested whether AlgK and AlgX are MucD substrates. Notably, degradation assays demonstrated that both the MucD 12-mer and trimer are able to completely degrade full-length AlgK and AlgX within two hours, with no significant difference in the degradation efficiency between the two oligomeric states (Fig. 1B, Fig. S1 C-E). Additionally, no partially degraded fragments were detected by SDS– PAGE, suggesting complete substrate turnover.

**Fig. 1.**
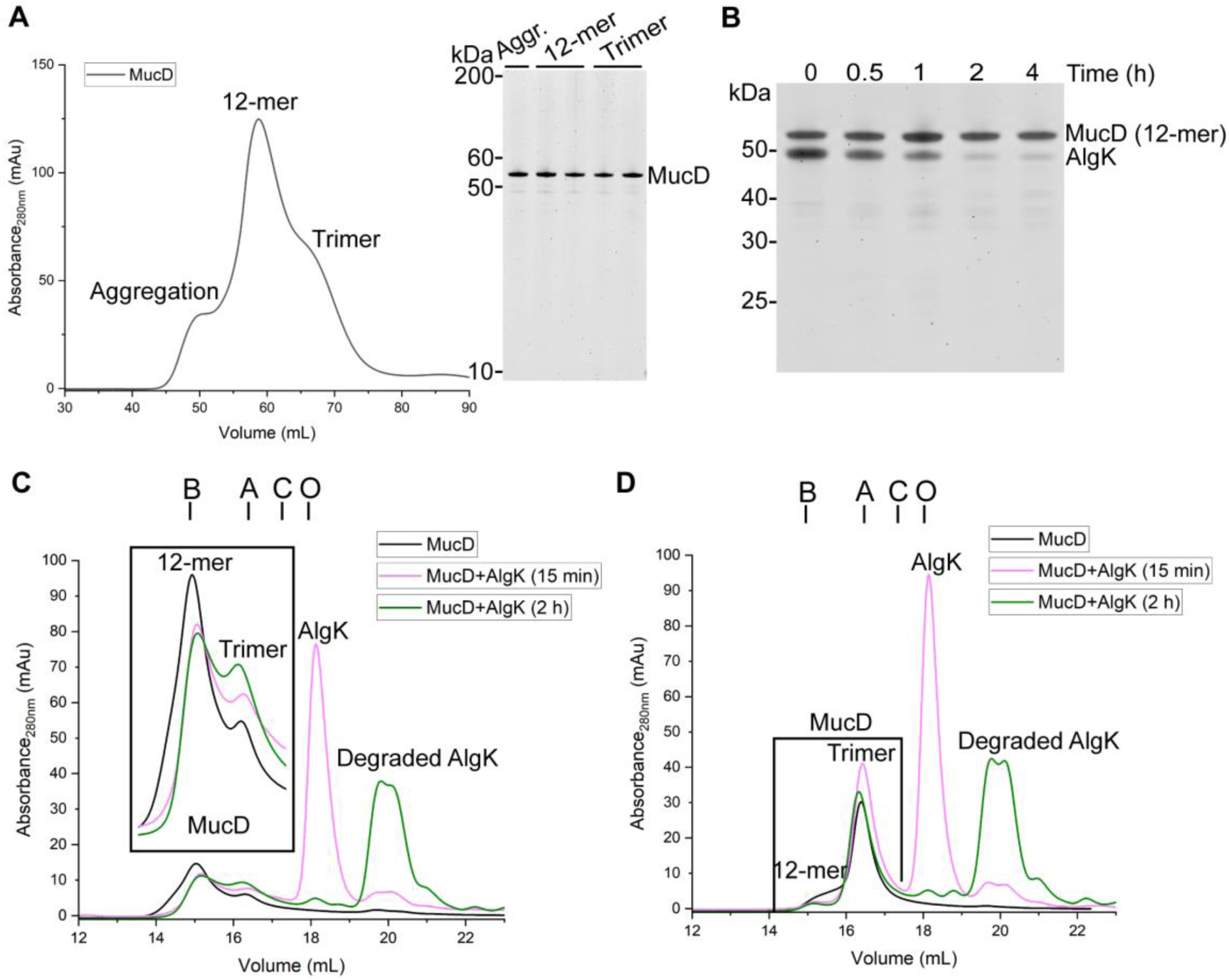
MucD oligomerization and proteolytic degradation of AlgK and AlgX. **(A)** The size-exclusion chromatography (SEC) profile of MucD (left panel). The purified MucD protein was loaded onto a HiLoad 16/600 Superdex 200 gel filtration column, and the absorbance of elution was recorded at 280 nm. Fractions corresponding to the indicated peaks were analyzed by SDS-PAGE (right panel). Aggr.: Aggregation. **(B)** Degradation of AlgK by MucD. The MucD 12-mer was incubated in a 1:1 molar ratio with purified AlgK at 37°C. At the indicated time points, aliquots were taken and analyzed by SDS-PAGE. **(C and D)** The effect of substrate AlgK on MucD oligomerization states’ equilibrium. MucD 12-mer **(C)** or trimer **(D)** was incubated with AlgK for 15 minutes or 2 hours, and the oligomerization states were analyzed by analytical SEC using a Superose 6 Increase 10/300 analytical gel filtration column. Note that the theoretical molecular weights of the MucD trimer and 12-mer are 159 kDa and 636 kDa, respectively. Elution volumes of molecular weight standards are indicated at the top by vertical lines: bovine thyroglobulin (B), aldolase (A), conalbumin (C), and ovalbumin (O) with molecular weights of 669, 158, 75, and 44 kDa, respectively.

Substrate binding converts DegP from a resting hexamer into active higher-order oligomers, including 12-mer and 24-mer assemblies [11]. We therefore asked whether MucD undergoes a similar substrate-dependent change. AlgK was incubated with either the MucD trimer or 12-mer for 15 minutes or 2 hours; under these conditions, AlgK was partially degraded at 15 minutes and fully degraded after 2 hours, respectively (Fig. 1B, Fig. S1C). Analytical SEC results showed that a brief 15-minute incubation of the MucD 12-mer with AlgK caused limited substrate degradation, accompanied by a modest decrease in the 12-mer population and an increase in the trimer population. This shift did not revert after prolonged incubation, even when degradation was complete (Fig. 1C). A similar oligomerization behavior was observed when AlgK was incubated with the MucD trimer (Fig. 1D). Taken together, these results suggest that substrate binding shifts the MucD oligomeric equilibrium toward the trimer, which appears to be the preferred state during proteolysis.

AlgK and AlgX form an AlgKX complex, which is essential for alginate biosynthesis *in vivo,* and the structure of this complex was recently determined in *P. putida* [19]. To test whether the AlgKX complex is also a proteolytic target of MucD, we first reconstituted the complex by co-incubating purified AlgK and AlgX proteins from *P. aeruginosa,* followed by analytical SEC analysis (Fig. S2A). The dominant peak corresponding to the AlgKX complex had an apparent MW of approximately 110 kDa, in line with the theoretical MW of 105.1 kDa for a 1:1 AlgK-AlgX heterodimer. Subsequently, our degradation assays revealed that the AlgKX complex was resistant to MucD proteolysis, with little degradation after 8 hours and substantial degradation only after 24 hours of incubation (Fig. S2B). In contrast, isolated AlgK and AlgX were completely degraded by MucD within two hours of incubation (Fig. 1B, Fig. S1 C-E). These data indicate that AlgK and AlgX are MucD substrates as individual proteins, whereas the assembled AlgKX complex is protected from MucD-dependent degradation.

### High-resolution analysis of MucD-dependent proteolytic products of AlgK and AlgX

To better understand how MucD recognizes and processes its substrate, we combined Liquid Chromatography-Mass Spectrometry (LC-MS) with bioinformatics analysis to identify MucD-dependent degradation products of AlgK and AlgX. Briefly, AlgK or AlgX was incubated with MucD for 8 hours to promote extensive proteolysis. The resultant degradation peptides were then separated from intact proteins using a 10-kDa molecular weight cutoff (MWCO) concentrator prior to LC-MS analysis. MS data were further processed using the “Utilities for Mass Spectrometry Analysis of Proteins” (UMSAP) software [23]. In total, we detected 381 peptides matching AlgK, corresponding to 90.1% sequence coverage, and 321 peptides matching AlgX, corresponding to 94.7% sequence coverage (Fig. S3A). The average peptide lengths were 13 and 14 amino acids for AlgK and AlgX, respectively. These results are consistent with previous degradation assays (Fig. 1B, Fig. S1 C-E) and confirm extensive MucD-mediated proteolysis of both substrates. The background peptides generation was negligible, as only 39 peptides matching AlgK (15.6% sequence coverage) and 38 peptides matching AlgX (14.3% sequence coverage) from substrate-only incubations were detected by LC-MS (Fig. S3A).

Proteolysis cleaves a peptide bond, generating two shorter fragments. Under standard protease nomenclature, the substrate residue at the N-terminal side of the scissile bond is designated as the P1 residue, while the residue at the C-terminal side is designated as the P1’ residue [24]. UMSAP also calculates the relative frequency of cuts (RFC) at each P1 residue, providing a quantitative measure of cleavage preference for a protease. For AlgK, 212 cleavage sites were mapped to the primary sequence, and 17 sites with RFC value greater than 9 were classified as high-RFC (hRFC) residues (Fig. S3B). Given that no experimental structure is currently available for *P. aeruginosa* AlgK, we mapped these hRFC residues onto an AlphaFold-predicted AlgK model. Only about half of the hRFC residues were surface exposed (Fig. 2A). Two clusters of hRFC residues (A100, R101, and G103; L315, G317, and L319) were located in close spatial proximity, which may reflect limits in assigning the exact N-or C-terminal ends of polypeptide fragments. AlgX showed a similar pattern when hRFC residues were mapped onto the AlgX structure (PDB: 4KNC) [22]. Among 210 detected cleavage sites, eight were classified as hRFC residues (Fig. S3C), and only five were surface exposed (Fig. 2B). Recent work on the HtrA protease HTRA1 showed that degradation of folded substrates can proceed sequentially: initial cleavage at surface-exposed sites induces local structural relaxation, followed by unfolding of early degradation products and exposure of buried cleavage sites [25]. MucD may use a similar mechanism to fully degrade native AlgK and AlgX, despite the burial of many cleavage sites. This model also helps explain why the AlgKX complex resists MucD proteolysis (Fig. S2B, Fig. S4A). Particularly, AlgK Q366 and AlgX V212 are hRFC residues located at the complex interface (Fig. S4 B and C). These residues are surface exposed in the isolated proteins and may participate in initial MucD cleavage events, but become buried upon AlgKX complex formation and are therefore shielded from proteolysis (Fig. 2 A and B, Fig. S4).

**Fig. 2.**
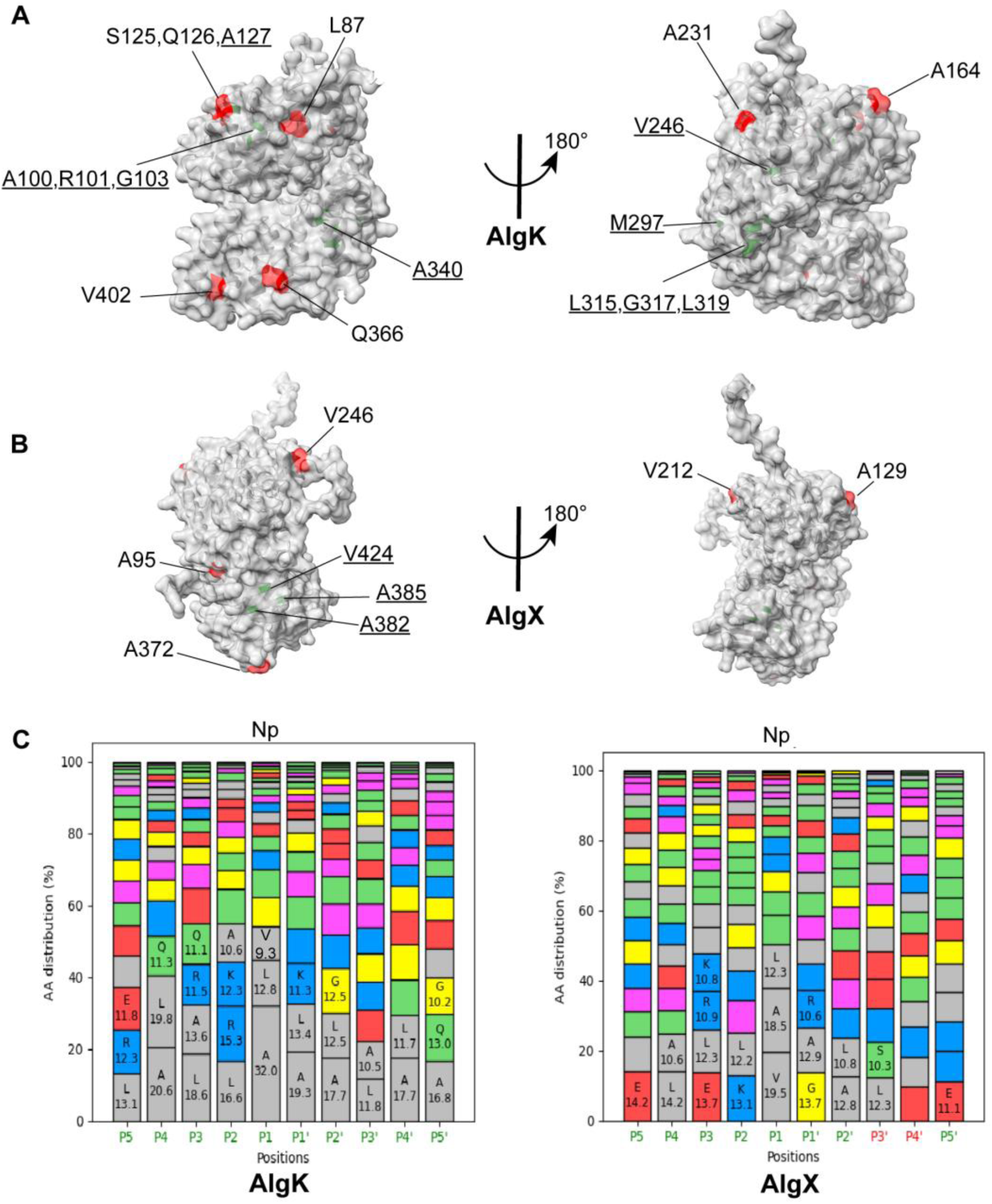
Analysis of the MucD-dependent proteolytic degradation products of AlgK and AlgX. AlgK **(A)** and AlgX **(B)** structural models with the high relative frequency of cuts (hRFC) residues highlighted. The AlphaFold-predicted AlgK (ID: AF-P96956-F1) and AlgX (PDB: 4KNC) models are shown as surfaces from two views rotated by 180°. Residues with RFC > 9 that are exposed to the surface (red) or buried inside (green; underlined) are highlighted and labelled. **(C)** Amino acid distribution of P5-P5’ sites for AlgK (left) and AlgX (right) degradation products. Amino acids with high distribution frequency at each site are labeled. “Np” at the top of the chart represents that non-polar amino acids are preferred at the P1 site. Colors for the P5-P5’ labels at the bottom of the chart: there is (green) or is not (red) a significant difference from the distribution expected under the assumption of no positional selectivity. The level of significance used for the χ2 test was 0.05. Colors of the bars represent amino acids with different chemical properties: gray for nonpolar residues (A, V, I, L, and M); green for polar amino acids (S, T, N, H, C, and Q); blue for positively charged amino acids (R and K); red for negatively charged amino acids (D and E); purple for aromatic amino acids (F, Y, and W), and yellow for special residues (G and P).

Previous studies suggested that DegP preferentially cleaves after hydrophobic P1 residues with short side chains and shows limited specificity across P2-P5 positions, which often contribute to substrate recognition and proteases binding [23,26]. We therefore analyzed MucD sequence specificity by calculating amino acid distributions from P5 to P5′ using UMSAP. Overall, MucD showed limited sequence specificity for either AlgK or AlgX (Fig. 2C), indicating that MucD acts as a protease that recognizes substrate conformation in addition to primary sequence. In particular, short side-chain non-polar residues, such as alanine, valine, and leucine, were enriched at P1 position and accounted for more than 50% of P1 residues in both AlgK and AlgX degradation products (Fig. 2C). Given that non-polar residues are frequently buried in folded proteins, this enrichment is consistent with our observation that approximately half of the hRFC residues in AlgK and AlgX are not surface exposed. It also supports a model in which initial cleavage exposes buried non-polar sites, enabling complete substrate degradation by MucD.

UMSAP also applies a χ2 test to determine whether the observed P5-P5’ residue distributions differ significantly from a theoretical null distribution representing a protease with no positional selectivity. This null distribution is primarily based on the overall amino acid abundance of the substrate [23]. For both AlgK and AlgX, the analysis revealed statistically significant selectivity across P5-P1 positions (Fig. 2C). For example, alanine, leucine, and valine were the three most enriched residues at the P1 site for both substrates. In contrast, P3′ and P4′ showed significant selectivity for AlgK but not for AlgX. Collectively, the χ2 test confirms the presence of specific, non-random amino acid patterns at the P5–P1 sites for both substrates. These distinct sequence profiles likely contribute to defining the substrate characteristics recognized by MucD.

### Structure of MucD in complex with its substrate polypeptide from AlgK

To better understand the oligomeric state of MucD and the structural basis of substrate recognition, we initially sought to determine the structure of wild-type MucD, but did not succeed despite extensive efforts. We therefore adopted a strategy commonly used for structural studies of proteases, including the HtrA-like protease DegP [27] and the AAA protease FtsH [28], by generating a catalytically inactive variant. Specifically, we purified MucD(S217A), in which the catalytic serine is replaced by alanine, as the structural target. The SEC profile of MucD(S217A) resembled that of wild-type MucD, with both 12-mer and trimer populations present (Fig. S5A). Protein degradation assays confirmed that MucD(S217A) is protease deficient, as neither AlgK nor AlgX was degraded after 4 hours of incubation (Fig. S5B), in contrast to the wild-type MucD (Fig. 1B, Fig. S1 C-E). Analytical SEC analysis further demonstrated that these two oligomeric states remain in dynamic equilibrium in solution (Fig. S5 C and D).

Interestingly, incubation of MucD(S217A) 12-mer with AlgK reduced the trimer population and promoted formation of 24-mer oligomers, which is distinct from that of the wild-type MucD (Fig. 1C, Fig. S5E). SDS-PAGE analysis of the 24-mer and 12-mer peaks showed co-elution of MucD with AlgK, suggesting that 24-mer formation is likely driven by substrate interaction (Fig. S5E). Next, we asked whether the interaction between MucD(S217A) and its substrates can be resolved by cryo-EM. To improve sample homogeneity for structural study, we examined AlgK degradation products and selected several polypeptides containing hRFC P1 residues as candidate substrate peptides (Fig. S3B). These peptides were chemically synthesized and tested their effect on the MucD(S217A) SEC profile. Eventually, we found that co-incubation of MucD(S217A) with AlgK_369-388_ (AlgK residues 369-388, N’-VDHLILAARAGQASADMALA) similarly induced 24-mer formation (Fig. S5F), indicating that AlgK_369-388_ binds to MucD. Cryo-EM analysis then yielded structures of the MucD(S217A) 12-mer and 24-mer at resolutions of 4.6 Å and 3.5 Å, respectively (Fig. S7 A and B, Table S2). The 12-mer and 24-mer complexes have overall diameters of approximately 155 Å and 190 Å, as well as enclosing an internal cavity of approximately 75 Å and 110 Å in diameter, respectively, dimensions comparable to the corresponding DegP oligomers [11,27] (Fig. 3A, Fig. S6A). Notably, both MucD(S217A) assemblies adopt relatively closed cage conformation, as the largest surface pores are only 8 Å in the 12-mer and 18 Å in the 24-mer, compared with 20 Å and 50 Å in the DegP 12-mer and 24-mer, respectively (Fig. S6A). Structural alignment of MucD and DegP 12-mers showed that the smaller pore in the MucD cage is mainly attributed to a shift of PDZ2* domains, where PDZ2* denotes a PDZ2 domain contributed by a neighboring trimer in inter-trimer interactions and oligomer assembly unless stated otherwise. This shift allows the “cage loops” (LC) to extend further toward the pore center (Fig. S6B). Consistently, sequence alignment showed that the MucD LC is six amino acids longer than that of DegP (Fig. S6C), allowing it to occupy more space and further restrict pore access. Superposition of the MucD 12-mer and 24-mer indicated that both assemblies use the same PDZ1-PDZ2*-mediated inter-trimer interaction, similar to what was previously reported for DegP 12-mer and 24-mer assemblies [27] (Fig. S6D). Thus, formation of distinct MucD oligomers appears to be enabled by conformational rearrangement of PDZ2 domains from neighboring trimers, facilitated by the flexible linker between the two PDZ domains within each protomer (Fig. S6D).

**Fig. 3.**
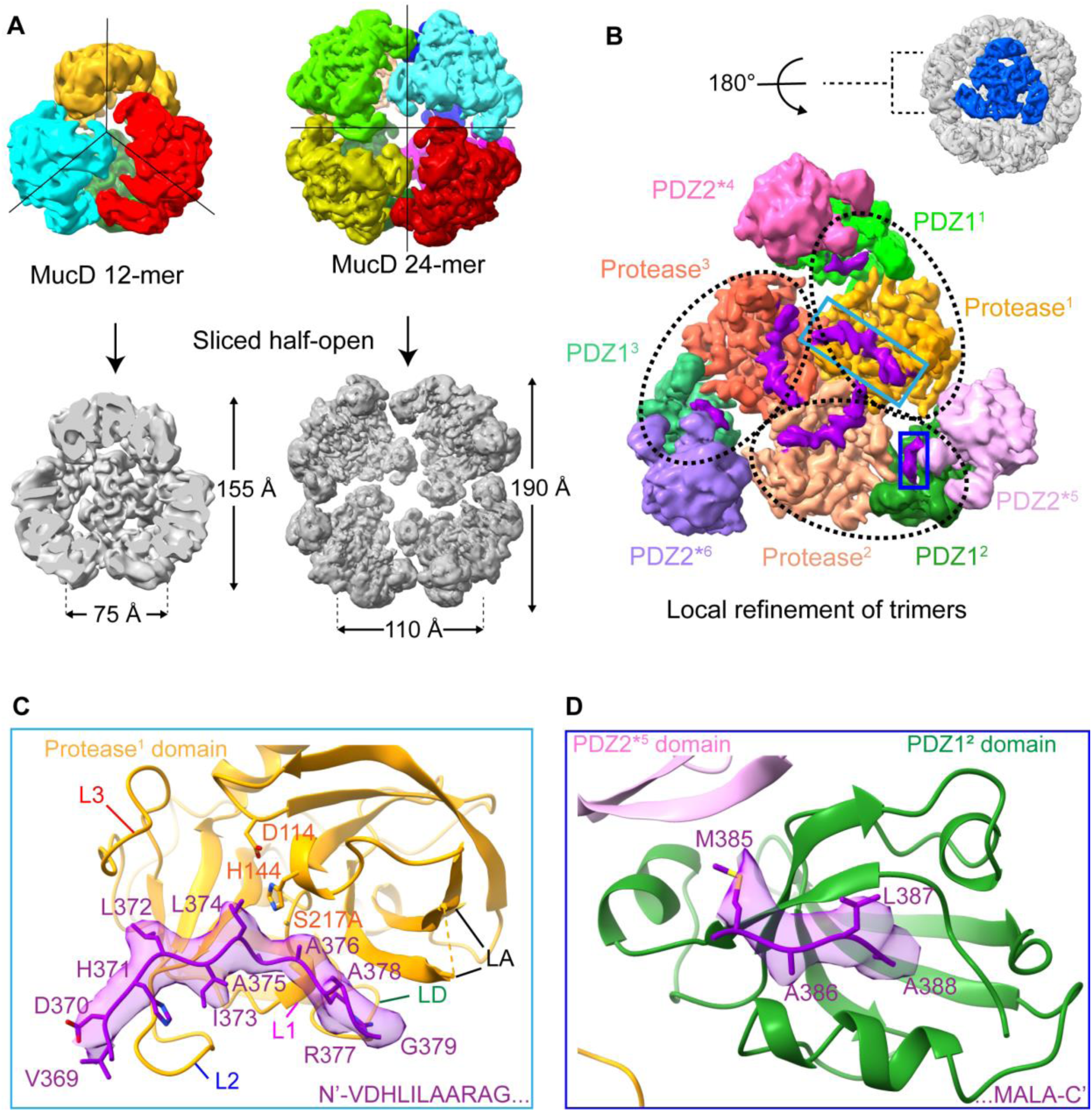
Structure of the MucD-AlgK_369-388_ complex. **(A)** Cryo-EM density maps of MucD 12-mer (PDB: 24XX) and 24-mer (PDB: 22KG). Upper panel: the overall architectures of the MucD 12-mer and 24-mer with tetrahedron or octahedron symmetry applied, respectively. Each trimeric subunit is shown in a different color. Lower panel: sliced half-open MucD 12-mer and 24-mer. The dimensions of the overall cage and the inner cavity are indicated. **(B)** Cryo-EM density map of the locally refined MucD trimers from the 24-mer (PDB: 22SK). Local refinement of an individual MucD trimeric face (the refinement mask is indicated in blue on the top right) within the 24-mer (gray) is shown (top right, view from “outside”). The protease and PDZ1 domains in each of the three protomers from a trimer are indicated by the superscript digits 1, 2, and 3 and outlined with dotted black ovals (lower panel, view from “inside”). Note that the PDZ2 domains of the protomers 1, 2, and 3 are not visible in this view. Protease: the protease domain; PDZ1: the PDZ1 domain; PDZ2*: the PDZ2* domains from different protomers are contributed by neighboring trimers as the superscript digits 4, 5, and 6 indicate. The extra density of the bound peptides is shown in purple. **(C)** A close-up view of the AlgK_369-388_ binding site in the protease domain of MucD. AlgK_369-388_ residues are shown in sticks overlaid with the corresponding density map. The catalytic triad residues (D114, H144, and S217A) are shown in sticks. The conserved L1, L2, L3, and LD loops of the protease^1^ domain, which are in catalytically active conformations, are indicated, respectively. The LA loop, which lacks well-resolved electron density, is connected with a dotted line. **(D)** A close-up view of the AlgK_369-388_ binding sites in the PDZ1^2^ domain of MucD. AlgK_369-388_ residues are shown in sticks overlaid with the corresponding density map.

To gain insight into the interaction interface, we generated locally refined maps of constituent MucD trimers at 2.9 Å and 3.0 Å resolution from the 12-mer and 24-mer reconstructions, respectively. This was achieved by symmetry expansion followed by local refinement using a mask covering one trimeric subunit (Table S2, Fig. S7 A and B). In both maps, additional density was clearly visible near the protease and PDZ1 domains of each protomer (Fig. 3B). Based on structural comparison with the DegP–hTRF1 (PDB: 8F0A) complex [27] (Fig. S8A), we assigned this density to the bound AlgK_369-388_ peptide. Structural alignment of the locally refined trimers from the 12-mer and 24-mer complexes showed that they are nearly identical, with a Cα RMSD of 0.43 Å (Fig. S8B). This similarity indicates that the MucD trimer serves as the basic building block of higher-order oligomers and the oligomeric state does not substantially alter the substrate-binding mode (Fig. S8B). Because the locally refined trimer from the 24-mer complex showed better overall map quality (Fig. S7 A and B), we used this model for subsequent structural analysis unless otherwise stated.

The fitted AlgK_369-388_ model revealed that the N-terminal 11 residues bind the protease domain of one protomer, whereas the last 4 residues of the C-terminus extend toward the PDZ1 domain of the adjacent protomer (Fig. 3 C and D). Density was well resolved for most of the bound AlgK_369-388_ peptide, with consecutive side chains clearly distinguishable, despite the fact that the residues Q380-D384 could not be modelled due to the poor electron density (Fig. 3 C and D). Notably, the MucD protease domain contains several conserved regulatory loops (LD, L1, L2, and L3) based on sequence and structure alignments with other bacterial HtrA-like proteases [11,20,28] (Fig. S9 A and B). Among these loops, LD, L1, and L3 regulate the switch between the active and inactive forms of the protease domain, while L2 determines the substrate specificity [11]. In the MucD structure, the protease domain adopts an open, catalytically competent conformation that accommodates the substrate peptide, as indicated by the displaced positions of these conserved regulatory loops that surround the bound peptide (Fig. 3C). Alignment of MucD and DegP L2 loops revealed differences in the composition of substrate-contacting residues, which likely underlies their distinct substrate specificities (Fig. S9 C and D). For example, R238 in MucD forms a salt bridge with D370 of AlgK_369-388,_ which significantly increases the capacity of the L2 loop to engage negatively charged substrate residues. Furthermore, F236 in MucD forms hydrophobic contact with L372 of the bound AlgK_369-388_ and may also support π-π interactions with aromatic residues in other substrates (Fig. S9C). Consistent with this role, substitution of F236 with alanine markedly reduced MucD proteolytical activity without disrupting oligomeric equilibrium: only limited AlgK degradation by MucD(F236A) was observed after 4 hours of incubation (Fig. S9 E and F, Fig. 1B, Fig. S1C). These results indicate that F236-mediated hydrophobic interaction is important for substrate recognition.

The side chain of the catalytic residue S217A points toward the peptide bond between substrate residues A375 and A376, defining the positions of the P1 and P1’ residues (Fig. 3C). Analysis of the peptide–protease interface showed that the P1 side chain, A375, inserts into a shallow hydrophobic S1 pocket formed primarily by these residues I212, P214, and I235 in MucD (Fig. S10 A and B). This arrangement is consistent with our MS analysis, which indicated a preference for small hydrophobic residues at the P1 position (Fig. 2C). Indeed, *in silico* substitution of A375 with seven other hydrophobic residues showed that alanine, valine, isoleucine, and leucine can be accommodated, whereas methionine, phenylalanine, tyrosine, and tryptophan, which have larger side chains, would exceed the spatial constraints of the S1 pocket and cause steric clashes (Fig. S10C). These structural observations agree well with the MS-derived specificity profile of the P1 site.

At the C-terminus of the AlgK_369-388_ peptide, only the last 4 residues showed sufficient electron density for modelling, suggesting a relatively weak but defined interaction with PDZ1 (Fig. 3D). The backbone carbonyl of substrate residue A388 forms hydrogen bonds with the backbone amides of MucD G273 and V274 (Fig. S11A). In addition, substrate residue L387 makes hydrophobic contact with MucD V275 through side-chain interaction, further stabilizing peptide binding (Fig. S11A). Notably, the side chain of A388 points into a hydrophobic pocket formed mainly by MucD residues L272, L331, P332, and V335 (Fig. S11 B and C). Because the PDZ1 domain functions as a central regulatory module in HtrA-like proteases, linking substrate recognition to protease activation [6], this hydrophobic PDZ1 pocket, together with adjacent residues V275 and L272, may recognize characteristic substrate C-terminal motifs and thereby promote allosteric activation of degradation. To test the role of PDZ1, we generated a MucD variant lacking this domain (MucD ΔPDZ1). This variant remained trimeric in solution (Fig. S11D), consistent with the observation that trimer formation is primarily mediated by protease-domain interactions (Fig. S8C). However, deletion of PDZ1 almost completely abolished proteolytic activity: no substantial degradation of AlgK or AlgX was detected after four hours of incubation (Fig. S11E), compared to the wild-type MucD (Fig. 1B, Fig. S1 C-E). These results indicate that substrate engagement by PDZ1 is essential for efficient MucD-mediated proteolysis.

### The proteolytic activity of MucD is regulated via oligomerization

Given that MucD exists in a dynamic equilibrium between trimeric and 12-mer states in solution, we next examined how oligomerization affects its proteolytic activity. Our structures show that higher-order MucD oligomers, including the 12-mer and 24-mer, are assembled through interactions between PDZ1 and PDZ2* domains (contributed by neighboring trimers) in an oligomerization mode reminiscent of DegP [27] (Fig. 4A, Fig. S6B). In DegP, formation of large oligomers is stabilized by a strong hydrophobic interface involving PDZ1 residues L276, M280, and F289 and PDZ2* residues Y431, Y444, and L446 [30]. By contrast, the corresponding MucD interface is dominated by a relatively weak π-π interaction between F287 (PDZ1) and F467 (PDZ2*) (Fig. 4A, right). Substitution of F467 with alanine yielded a variant that remained trimeric after purification, as shown by analytic SEC, and no longer underwent substrate-induced changes in oligomeric state (Fig. 4B). These structure-guided mutagenesis results support a model in which PDZ1-PDZ2* interactions drive higher-order MucD assembly. Notably, MucD(F467A) degraded AlgK more efficiently than the wild-type MucD, accomplishing degradation of full-length AlgK within one hour (Fig. 4D). However, SDS-PAGE in combination with MALDI/TOF-MS analysis revealed accumulation of partially degraded AlgK fragments (Fig. 4D, Table S3), unlike wild-type MucD, which degraded AlgK completely without detectable large fragment bands (Fig. 1B, Fig. S1C).

**Fig. 4.**
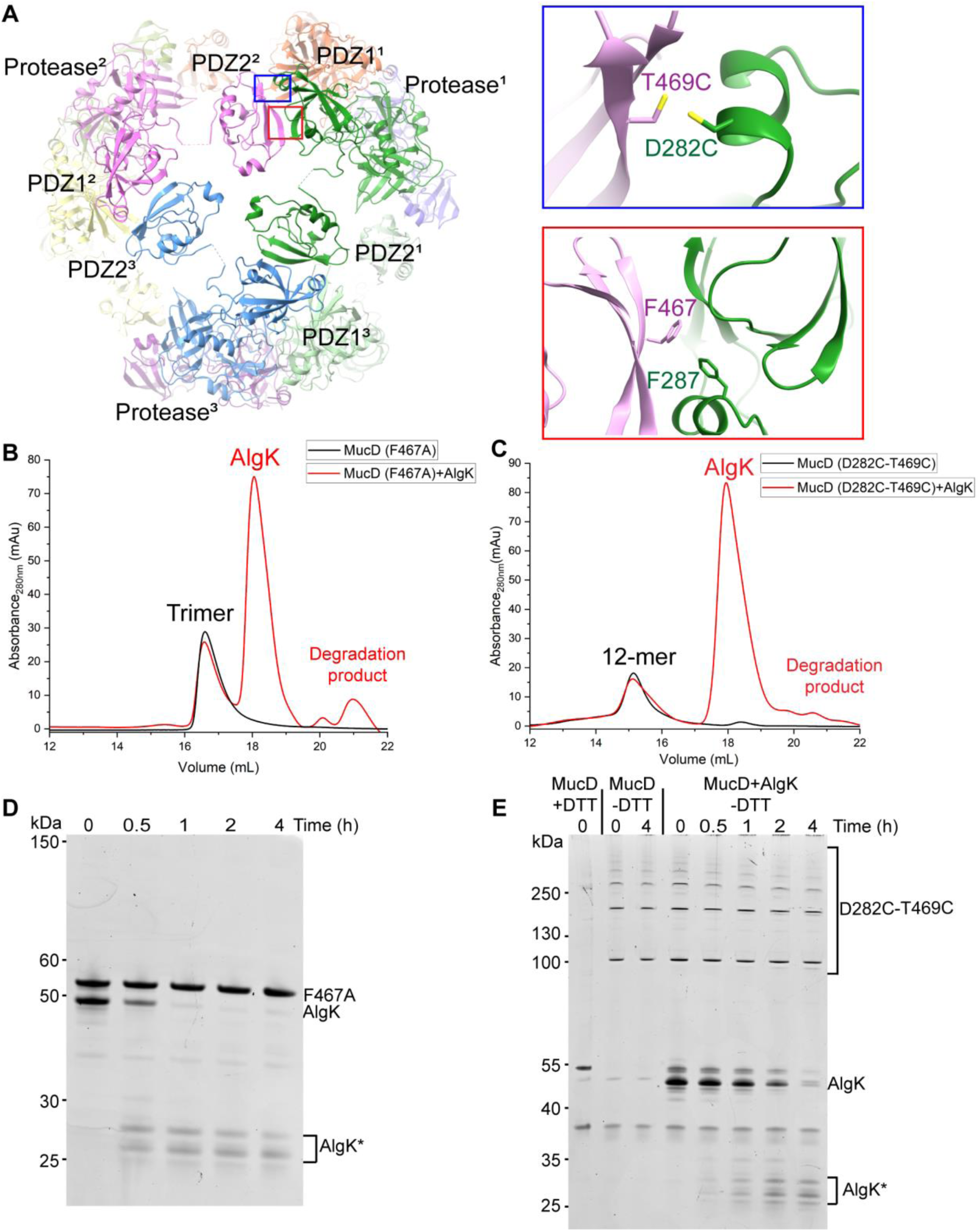
Regulation of MucD proteolytic activity through oligomerization. **(A)** Characterization of MucD inter-trimer interactions. MucD 12-mer (PDB: 24XX) is shown in cartoons with each protomer colored the same and indicated with superscript digits 1, 2 and 3. The flexible linker between the PDZ1 and PDZ2 domains is shown in dotted lines. Red rectangle: close-up view of interacting residues F287 and F467 (shown in sticks) at the PDZ1-PDZ2* domains’ interface. Blue rectangle: close-up view of residues D282C and T469C targeted for cysteine replacement. **(B and C)** MucD (F467A) **(B)** and MucD (D282C-T469C) **(C)** were incubated in a 1:1 molar ratio with AlgK for 15 min. The oligomeric states were analyzed by analytical SEC using a Superose 6 Increase 10/300 GL gel filtration column. MucD mutants without substrate addition served as a control. **(D and E)** Degradation of AlgK by MucD mutants. MucD(F467A) **(D)** and MucD(D282C-T469C) **(E)** were incubated with AlgK in a 1:1 molar ratio at 37°C. At the indicated time points, aliquots were taken and analyzed by SDS-PAGE. AlgK*: partially degraded AlgK identified by MALDI/TOF-MS. +/-DTT: with or without 20 mM dithiothreitol (DTT) in the SDS-loading dye.

We propose that the MucD 12-mer adopts a closed cage conformation that restricts substrate access, thereby limiting its proteolytic activity against folded substrates (Fig. S6 A and B). To test this model, we introduced cysteine substitutions at D282 and T469 to reinforce the PDZ1– PDZ2* interface, given that both residues are solvent-exposed at the interface and have side-chain lengths comparable to cysteine (Fig. 4A). As MucD is a periplasmic protein with only one endogenous cysteine within its signal peptide, the non-reducing periplasmic environment should favor formation of specific disulfide bonds at the engineered sites [30]. Indeed, the purified MucD(D282C–T469C) mutant predominantly exists as a stable 12-mer, with no tendency to disassemble upon substrate addition (Fig. 4C). SDS-PAGE analysis revealed prominent DTT-sensitive high-molecular-weight bands, indicating successful interdomain cross-linking via disulfide bonds (Fig. 4E). If cross-linking occurred at every D282C-T469C pair across the PDZ1-PDZ2* interface within the 12-mer, non-reducing SDS-PAGE would be expected to show a band of ∼160 kDa, corresponding to three disulfide-linked MucD molecules from adjacent trimers (Fig. 4A). Of note, additional bands near 110 kDa likely reflect incomplete cross-linking across the interfaces, whereas minor bands above 250 kDa may represent nonspecific aggregation caused by random cross-linking *in vivo* (Fig. 4E). Functionally, MucD(D282C–T469C) degraded AlgK much more slowly than wild-type MucD (Fig. 1B, Fig. S1C), with both full-length and partially degraded AlgK still detectable after 4 hours of reaction (Fig. 4E), supporting the hypothesis that the 12-mer is less proteolytically active. Although cage assembly does not fully abolish proteolysis, the dynamic equilibrium between the 12-mer and trimer likely enables complete substrate degradation while preventing accumulation of partially degraded fragments, thereby supporting periplasmic protein quality control.

Within each MucD trimer, protomer association is stabilized by extensive hydrophobic contacts and hydrogen bonds between protease-domain residues at the interface (Fig. S8C), similar to the trimeric interface observed in *P. syringae* MucD (psMucD; PDB: 8K2Y) [20]. At the N-terminal region of MucD, where the three protomers converge, F30 and W171 form a triangular aromatic arrangement stabilized by edge-to-face or stacked π-π interactions (Fig. S8C). Additional π-π interactions occur between F236 from one protomer and F179–F181 from a neighboring protomer (Fig. S8C). Hydrogen bonds between the amide groups of Q207 and R195 from one protomer and the hydroxyl group of S184 from another protomer further reinforce the trimer stability (Fig. S8C). Structural alignment showed that trimeric psMucD (PDB: 8K2Y) and *P. aeruginosa* MucD (paMucD; PDB: 22SK, from the 24-mer reconstruction) are highly similar, with a Cα RMSD of 0.69 Å (Fig. S8B). In both structures, the major regulatory loops of the protease domain, including L1, L2, L3, and LD, adopt open conformations that allow substrate binding (Fig. 3C, Fig. S9B). These observations support the view that trimeric MucD represents an active state for proteolysis.

### MucD represses alginate production by directly degrading alginate synthesis proteins

MucD has been reported as a negative regulator of alginate production in *P. aeruginosa*, as the deletion of *mucD* leads to alginate overproduction in the wild-type laboratory PAO1 strain [14,18]. To determine whether this regulation depends on MucD protease activity, we measured alginate-producing levels in Δ*mucD* mutants in the PAO1 background using the carbazole assay, a quantitative method for detecting alginate in *P. aeruginosa* [19,32]. Notably, alginate production was below the detection limit in wild-type PAO1 but increased significantly in the Δ*mucD* mutants (Fig. S12A), in line with the previous reports [14,33]. Complementation with wild-type *mucD* or proteolytically active MucD(F467A) variant expressed from the commonly used plasmid pUCP24 [33] reduced alginate production to an undetectable level (Fig. S12A). Growth monitoring over 24 hours confirmed that these differences were not caused by changes in cell fitness (Fig. S12B). In contrast, complementation with either the proteolytically inactive MucD(S217A) variant or the disulfide-locked 12-mer MucD(D282C-T469C) variant did not significantly reduce alginate production (Fig. S12A). These results indicate that MucD-mediated repression of alginate production requires proteolytic activity and is impaired when MucD is trapped in the resting 12-mer state.

A previous study reported that deletion of *mucD* in PAO1 increases promoter activity of the *alg* operon [14]. To assess whether the elevated alginate production observed here was accompanied by transcriptional changes in *algK and algX*, quantitative reverse transcription PCR (RT-qPCR) for the wild-type, Δ*mucD,* and Δ*mucD*::*mucD* strains was carried out. The results showed that the mRNA levels of *algK* and *algX* did not differ significantly among the strains. By contrast, *algD*, the proximal gene in the operon encoding the rate-limiting enzyme of the pathway, increased ∼2.8-fold in the Δ*mucD* strain and restored to wild-type levels upon *mucD* complementation (Fig. S12C). The different behavior of the proximal *algD* transcript and the distal *algK* and *algX* transcripts likely reflects natural transcriptional attenuation or differential mRNA stability across this large 12-gene biosynthetic cluster, consistent with previous evidence for increased promoter activity in Δ*mucD* cells [14,35,36]. Given that AlgK is a MucD substrate *in vitro*, we next examined the AlgK protein level *in vivo* by Western blot in PAO1, Δ*mucD*, and Δ*mucD*::*mucD* strains. Our results revealed that AlgK protein level was markedly increased in the Δ*mucD* mutant and restored by *mucD* complementation (Fig. S12D). Together with its degradation by MucD *in vitro* (Fig. 1B, Fig. S1C) and its reported association with MucD *in vivo* [14], these findings support AlgK as a physiological substrate of MucD. AlgX can also be degraded by MucD *in vitro* (Fig. S1 D and E) and was previously identified in the same periplasmic complex, however, its substrate status *in vivo* requires more experiments with complementary approaches.

Previous studies demonstrated that alginate export and modification require a large complex composed of AlgK, AlgX, AlgJ, and AlgF in *P. aeruginosa* [37]. To test whether MucD also targets other alginate synthesis proteins in this complex, we purified soluble constructs of AlgJ_79-379_ and AlgF_30-216_ [37,38] and examined their susceptibility to MucD-mediated degradation. Interestingly, AlgF was degraded by MucD, whereas AlgJ was resistant (Fig. S13 A and B), suggesting that post-translational regulation by MucD is selective and restricted to particular components of the alginate biosynthetic machinery. Collectively, these results suggest that MucD can alternatively regulate alginate production by directly degrading alginate biosynthetic proteins.

## Discussion

MucD is an HtrA-like serine protease that plays a critical role in stress adaptation, virulence, and regulation of alginate biosynthesis in *P. aeruginosa* [16,18]. In this study, we newly identified AlgK and AlgX as MucD substrates. Both proteins are essential components of the alginate biosynthetic machinery, responsible for alginate chain export and acetylation, respectively. By combining LC-MS with *in silico* mapping of the proteolytic fragments using UMSAP, we defined MucD cleavage preferences and sequence selectivity. Notably, our cryo-EM structures of the MucD 12-mer and 24-mer complexed with a substrate peptide uncover the structural basis for MucD oligomerization, protease-substrate interactions, and specificity, which nicely rationalizes our biochemical characterization data. Mutagenesis of interface residues demonstrated that the proteolytic activity of MucD is modulated by its oligomeric state. Finally, we showed that beyond the known transcriptional-level regulation, MucD alternatively represses alginate production *in vivo* through degradation of alginate biosynthesis proteins. These findings establish a post-translational regulatory mechanism for alginate biosynthesis and provide mechanistic insight into the function of bacterial HtrA-like proteases.

### Oligomerization regulates MucD proteolytic activity

Oligomerization is widely employed for HtrA-like proteases to regulate their activity [6]. DegP, for example, forms a proteolytically inactive hexamer through the interaction between the extended LA loops of two “face-to-face” trimers [38]. Upon substrate binding, DegP rapidly converts into higher-order oligomers (12-mer and 24-mer), driven by hydrophobic interactions between PDZ1 and PDZ2* domains, and then returns to the resting state after substrate degradation [12]. These cage-like assemblies have also been proposed to support chaperone activity by assisting the folding of outer membrane proteins [11]. However, their physiological significance remains debated, as higher-order oligomers are not strictly required for proteolysis or high-temperature survival [29].

In contrast, MucD follows a distinct regulatory pattern, namely maintaining an equilibrium between trimeric and 12-mer states in solution (Fig. 1A, Fig. S1 A and B). Substrate binding promotes the formation of the trimer and disassembly of the 12-mer, suggesting that the trimer is the preferred active form (Fig. 1 C and D). Mutagenesis of inter-trimer interface residues further supports the notion that the 12-mer represents a resting or less active state, while the trimer mediates efficient proteolysis (Fig. 4). Nevertheless, none of the MucD variants tested fully reproduced wild-type substrate turnover, and both oligomer-biased mutants accumulated partially cleaved AlgK fragments (Fig. 4 D and E). These observations suggest that dynamic exchange between the 12-mer and trimer may be important for processive degradation, perhaps by facilitating substrate remodeling or unfolding to expose buried cleavage sites.

The distinct oligomerization-driven regulation of MucD and DegP is structurally relevant. While both enzymes use a similar trimeric building block, MucD has a shorter LA loop than DegP, which may limit its ability to form a stable hexamer (Fig. S9A). Furthermore, the PDZ1-PDZ2* interface that sustains the MucD 12-mer or 24-mer cage is mediated mainly by a π-π interaction between F287 and F467 (Fig. 4A). This interface is substantially weaker than the multi-residue hydrophobic interface in DegP, where PDZ1 residues L276, M280, and F289 interact with PDZ2* residues Y431, Y444, and L446 [29]. These differences likely explain why MucD and DegP show distinct patterns of cage assembly and disassembly during proteolysis (Fig. 1 C and D).

Interestingly, substrate addition to the proteolytically inactive MucD(S217A) mutant shifted the equilibrium toward 12-mer and 24-mer assemblies, a behavior distinct from that of wild-type MucD (Fig. 1 C and D, Fig. S5 E and F). This finding raises the possibility that inhibitory peptides or small proteins, analogous to naturally produced serine protease inhibitors such as serpins [39], could regulate MucD by stabilizing the higher-order oligomeric states as post-cleavage intermediates. Ecotin, a broad-spectrum periplasmic serine protease inhibitor found in many bacteria, including *E. coli* and *P. aeruginosa*, provides one example of this type of post-translational regulation [40]. Although we did not observe 24-mer formation for wild-type MucD, the 12-mer and 24-mer cage assemblies may also relate to a potential chaperone-like function. Under stress conditions, membrane or periplasmic proteins with critical function may be recognized by MucD and trigger the cage formation. Since the higher oligomers of MucD adopt a more closed formation (Fig. S6A), they may shield and stabilize captured substrates from the disordered periplasm environment. However, we did not detect proteins co-purifying with MucD, and no putative chaperone partners have yet been reported. Given that the chaperone activity is highly condition-dependent, whether MucD has chaperone activity remains an open question.

### Substrate recognition and specificity

Hydrophobic residues are preferred cleavage sites for many HtrA-like proteases [8,41]. Consistent with this feature, structural analysis of the MucD S1 pocket together with MS-based mapping of cleavage products demonstrated that MucD preferentially cleaves at hydrophobic residues (Fig. 2C, Fig. S10). Substrate recognition, however, is not determined by the protease domain alone. Our structure of the MucD-AlgK_369-388_ complex revealed that PDZ1 also engages the substrate C-terminus through backbone hydrogen bonds and hydrophobic contacts, particularly with the side chain of the C-terminal end A388 positioned in a hydrophobic pocket analogous to the S1 site (Fig. S10, Fig. S11). In the complex structure (Fig. 3 C and D), the 13-residue peptide spanning from the P1’ residue A376 to the C-terminal end residue A388 closely matches the average length of AlgK and AlgX degradation fragments, which are 13 and 14 residues, respectively (Fig. S3A). Together with the observation that the deletion of the PDZ1 domain almost completely abolishes proteolytic activity, these findings support a model in which PDZ1 acts as a receptor for the substrate C-terminus following an initial cleavage event. This interaction may stabilize the protease-substrate complex and help define fragment length during processive degradation.

### Regulation of alginate biosynthesis and physiological implications

MucD has long been recognized as a negative regulator of alginate biosynthesis. Previous studies have suggested a transcriptional regulation function for MucD, as deletion of *mucD* increases promoter activity of the *alg* operon [14], consistent with our observation that *algD* mRNA levels were significantly elevated in the Δ*mucD* strain (Fig. S12C). Our results further demonstrate that MucD proteolytic activity is required for this regulatory function in wild-type *P. aeruginosa*. Notably, transcript levels of the essential alginate biosynthesis genes, *algK* and *algX,* remained unchanged in Δ*mucD* cells, whereas AlgK protein abundance increased significantly (Fig. S12 C and D). In large bacterial operons such as the *alg* operon, polycistronic transcripts can undergo progressive downstream attenuation, internal termination, or segmented mRNA decay, uncoupling steady-state transcript levels of distal structural genes (*algK* and *algX*) from primary promoter activity [34]. Collectively, these findings indicate that MucD regulates alginate biosynthesis through both negative transcriptional control of the *alg* operon and post-translational quality control of key assembly components, including AlgK and AlgX, as a model is proposed to demonstrate MucD oligomerization assembly, its activity regulation (Fig. S14A), and its involvement within the alginate biosynthesis multiprotein complex (Fig. S14B).

MucD is a periplasmic HtrA-like serine protease that involves pathways critical for bacterial survival and virulence, particularly for *P. aeruginosa,* an opportunistic pathogen capable of infecting plants, animals, and even humans. Loss of *mucD* reduces virulence in several infection models [16,17], underscoring its physiological importance. While our study elucidates a role for MucD in post-translational control of alginate biosynthesis, its broader involvement in other processes—such as host-pathogen interactions, antibiotic resistance, and biofilm formation—remains to be explored.

## Methods

### Bacteria strains and plasmids

All recombinant plasmids were generated by Gibson assembly using the ClonExpress II One Step Cloning Kit (Vazyme). The *mucD* gene variants encoding MucD S217A, F467A and D282C-T469C mutants were generated by site-directed mutagenesis. Seamless deletion of *mucD* gene in PAO1 strain was conducted using the pEX18Tc plasmid. Details of bacterial strains and plasmids used in this study are listed in Table S4.

### Protein purification

AlgK, AlgX as well as MucD and its variants from *P. aeruginosa* were purified similarly. Briefly, the plasmid pMMB67EH containing the relevant gene with a hexa-histidine (6×His) tag at the C-terminus was transformed into the *P. aeruginosa* PAO1 strain by electroporation for protein expression. Bacteria were cultured at 37°C in Luria-Bertani (LB) broth (10 g/L tryptone, 5 g/L yeast extract, and 5 g/L NaCl, pH 7.0-7.5) supplied with 150 μg/mL carbenicillin to OD_600nm_ reached 0.8 with agitation at 200 rpm. Proteins expression was induced by 0.8 mM isopropyl-β-d-thiogalactopyranoside (IPTG), and bacteria were grown at 16°C for 16 h with agitation at 160 rpm. Bacteria were harvested by centrifugation (4000 g, 20 min) and resuspended in Buffer A (50 mM HEPES and 500 mM NaCl, pH 7.5). Cells were lysed by sonication and then centrifuged at 20000g for 1 h to remove insoluble fractions. The supernatant was collected and incubated with nickel–nitrilotriacetic acid (NTA) beads (Cytiva), pre-equilibrated with Buffer A, at 4°C for 1 h with gentle rotation. The proteins were purified by gravity flow chromatography by extensive washing with Buffer E (20 mM HEPES and 150 mM NaCl, pH 7.5) supplemented with 30 mM imidazole and subsequent elution with Buffer E supplemented with 300 mM imidazole. Eluted proteins were concentrated using a 30-kDa molecular weight cutoff (MWCO) concentrator (Millipore) and loaded onto a HiLoad 16/600 Superdex 200 column (Cytiva) pre-equilibrated with Buffer E. The fractions containing the target proteins were pooled, concentrated, and stored by snap freezing for downstream experiments.

AlgJ_79-391_ and AlgF_30-216_ were cloned from the *P. aeruginosa* genome into the pDBHT-2 vector with a 6×His tag at the N-terminus, separately. The recombinant vectors were transformed into the *E. coli* BL21(DE3) strain for protein expression and purification as described previously [36,37]. Purified AlgJ_79-391_ and AlgF_30-216_ proteins were buffer exchanged to Buffer E, concentrated, and stored by snap freezing for downstream experiments.

### *In vitro* protein degradation assays

80 μM MucD and 80 μM substrates (AlgK, AlgX, AlgKX, AlgJ_79-391_ and AlgF_30-216_) were mixed in Buffer E and incubated at 37°C. At each time point, an aliquot was taken and diluted 400-fold with Buffer E. 50 ng of samples supplied with NuPAGE LDS Sample Buffer (Invitrogen) were incubated at 70°C for 10 min before loading onto a 4–12% NuPAGE Bis-Tris Mini Protein Gel (Invitrogen). Gel electrophoresis was conducted in MES buffer at 200V for 38 min. After electrophoresis, the gel was stained by SYPRO Ruby (Thermo-Fisher Scientific) and visualized by the ChemiDoc Touch Image System (Bio-Rad). Degradation assays of MucD variants were conducted following a similar process.

### Analytical Size Exclusion Chromatography (SEC)

The purified 12-mer and trimer population of wild type MucD and MucD(S217A) at a concentration of 120 μM were loaded to an analytical SEC column (Superose 6 Increase 10/300 GL, Cytiva). To analyze the proteolysis process or form complex, MucD and its variants were incubated with AlgK at a 1:1 molar ratio (120 μM) at 37 °C for 15 min or 2 h prior to application to the analytical SEC column (Superose 6 Increase 10/300 GL, Cytiva). Apparent molecular weights (MW) of different MucD oligomeric states were calculated by interpolating from MW *vs.* elution volume standard curve, which was generated by using a Gel Filtration Standard (Bio-Rad) as per the manufacturer’s guidelines.

### Analysis of MucD proteolysis products by LC-MS

For proteolysis, 80 μM MucD was incubated with 80 μM AlgK or AlgX in Buffer E at 37°C for 8 h. Samples were then centrifuged using a 10-kDa molecular weight cutoff (MWCO) concentrator (Millipore) to remove intact proteins. Peptides resulting from AlgK and AlgX degradation in the flowthrough were subjected to LC-MS analysis.

The mass spectrometry experiments were performed in the Mass Spectrometry Core Facility in the School of Biological Sciences, Nanyang Technological University. The fractionated peptides were separated and analyzed using a Vanquish Neo UHPLC System coupled to an Orbitrap Exploris 480 (ThermoFisher Scientific). Separation was performed on an EASY-Spray 75 μm × 15 cm column packed with PepMap Neo C18 2 μm, 100 Å (ThermoFisher Scientific) using solvent A (0.1% formic acid) and solvent B (0.1% formic acid in 80% ACN) at a flow rate of 300 nL/min with a 60 min gradient. Peptides were then analyzed on an Orbitrap Exploris 480 apparatus with an EASY nanospray source (ThermoFisher Scientific) at an electrospray potential of 2.0 kV. A full MS scan (200–2,000 m/z range) was acquired at a resolution of 60,000. Dynamic exclusion was set as 30 s. The resolution of the higher energy collisional dissociation (HCD) spectra was set to 30,000. The automatic gain control (AGC) settings of the full MS scan and the MS2 scan were 200% normalized AGC target and standard, respectively. The data-dependent mode was cycle time. The time between each master scan was 2 seconds. An isolation window of 1.6 m/z was used for MS2. Single and unassigned charged ions were excluded from MS/MS. For HCD, the normalized collision energy was set to 32%. Raw data files were processed and searched using Proteome Discoverer 2.2 (ThermoFisher Scientific). The raw LC-MS/MS data files were loaded into Spectrum Files (default parameters set in Spectrum Selector). The Mascot algorithm was then used for data searching to identify proteins using the following parameters: missed cleavage of three; dynamic modifications were oxidation (+15.995 Da) (M), and deamidation (+0.984 Da) (N and Q), acetylation (+42.0105 Da) (K), methylation (14.0157 Da) (K andR), (28.0313 Da) (K and R) and phosphorylation (+79.966 Da) (S, T, and Y). Percolator was applied to filter out the false MS2 assignments at a strict false discovery rate of 1% and a relaxed false discovery rate of 5%.

### Analysis of detected peptides by UMSAP

Identified peptides were further analyzed using the Targeted Proteolysis module of UMSAP 2.3.3 [23]. The significance level was set to 0.05 and the minimum score value to 10. A Log_2_ transformation was applied to the data before the analysis. The amino acid distribution around the cleavage sites included five residues in each direction. Calculation of the relative frequency of cuts by UMSAP followed a similar procedure as described in the reference [25].

### Multiple and pairwise sequence alignment

Protein sequences were retracted from UniProt [42]. The UniProt entry identifiers for the sequences are as follows: paMucD (G3XD20); psMucD (Q9AQD1); ecDegP (P0C0V0); ecDegQ (P39099). For both multiple and pairwise sequence alignment, sequences were input into Snapgene software.

### Cryo-EM samples preparation and data collection

The AlgK_369-388_ peptide (synthesized by GeneScript) was dissolved in 100% DMSO at the concentration of 2.4 mM. To form the MucD(S217A)-AlgK_369-388_ complex, AlgK_369-388_ was diluted by Buffer S (20 mM HEPES and 50 mM NaCl, pH 7.5) to 120 μM (0.5% DMSO) and incubated with 24 μM MucD(S217A) in Buffer S at 37°C for 16 h. The sample was loaded onto a Superose 6 Increase 10/300 gel filtration column (Cytiva) pre-equilibrated with Buffer S. The fractions containing the 12-mer or 24-mer population were pooled and concentrated to 5.0 mg/mL.

For both complexes, a total of 4 μl of protein sample at a concentration of 5.0 mg/ml was applied to the Quantifoil Cu R1.2/1.3 grids. The grids were blotted using a Vitrobot Mark IV (Thermo-Fisher Scientific) operated at 4°C and 100% humidity with a blotting time of 1 s, blotting force of 1, and waiting time of 1 s. Datasets were collected on a Titan Krios (Thermo-Fisher Scientific) 300-kV electron microscope equipped with a GIF Quantum energy filter and a Falcon4i detector with a defocus range between −0.6 μm and −1.2 μm. The magnification was ×130,000 with a pixel size of 0.97 Å.

### Cryo-EM data processing

Micrographs were imported to Cryosparc4.1 for patch motion correction and patch CTF estimation [43]. Micrographs with an estimated CTF resolution higher than 6 Å were selected for subsequent processing. Particles were auto-picked by the blob picker, manually curated, and extracted with a box size of 360 and a pixel size of 3.04 Å. All extracted particles were subjected to a reference-free 2D classification. Good classes with clear particle features were selected and used for initial model building and heterogeneous refinement. Only the reasonable models were selected, and the good particles were re-extracted with a box size of 320 and a pixel size of 1.216 Å. The re-extracted particles were used for homogeneous refinement and, subsequently, local refinement. The final resolutions of all resulting maps were calculated based on gold standard Fourier shell correlation (FSC) = 0.143. AlphaFold–predicted models of MucD (database accession code: AF-G3XD20-F1) were used for model building [44], which was manually adjusted with COOT [45], and real-space refinement was done in Phenix Suite [46]. All the structure-related figures were generated by ChimeraX [47].

### Alginate production, purification, and detection

Alginate production and purification were carried out as described previously [19] with several changes. Briefly, an overnight bacterial culture was diluted in 10 mL of alginate-producing medium (100 mM monosodium glutamate, 7.5 mM NaH_2_PO_4_, 16.8 mM K_2_HPO_4_, and 10 mM MgSO_4_, pH=7.0) to an OD_600nm_ of 0.04 and incubated at 37°C for 24 h with agitation at 200 rpm. For culturing *mucD* complementary strains, the medium was supplemented with gentamicin at a final concentration of 50 µg/mL. Cells were removed by centrifugation (4000 g, 15 min) and the supernatant containing alginate was collected. To precipitate alginate, three volumes of ice-cold absolute ethanol was added to the supernatant and incubated at 4 °C for 2 h with gentle rotation. Precipitated alginate was collected by centrifugation at 4 °C (4000 g, 20 min) and washed with 10 mL ice-cold absolute ethanol twice. The excess ethanol was removed by air drying samples at 25°C for 6 h. 5 mL of distilled water was added to dissolve alginate by thorough vortex and subsequent incubation at 42°C for 30 min.

Alginate was quantitatively detected using the carbazole assay for uronic acids as described previously [19,31] with several modifications. Briefly, a borate-sulfuric acid solution (0.1 M H_3_BO_3_ in concentrated H_2_SO_4_) and a carbazole reagent (0.1% (w/v) of carbazole in absolute ethanol) were prepared. 3 mL of the borate-sulfuric acid solution was chilled on ice. A 500 μL aliquot of alginate solution was slowly added to the top of the acid mixture. After a brief vortex at low amplitude, 100 μL of the carbazole reagent was added to the acid sample mixture on ice. Following a brief vortex, samples were incubated at 55 °C for 30 min and subsequently cooled at room temperature for five minutes. The absorbance was measured at 530 nm using a spectrophotometer with three biological repeats.

### RNA extraction, reverse transcription, and quantitative PCR

The overnight culture was diluted in 3 mL of alginate-producing medium to an initial OD_600nm_ of 0.04 and incubated at 37 °C for 24 h with agitation at 200 rpm. After harvesting by centrifugation, bacteria were normalized to an OD_600nm_ of 1.0 in 1 mL PBS buffer. Total RNA was isolated using a PureLink RNA Mini Kit (Invitrogen) and RNA concentrations were determined using a NanoDrop spectrophotometer (Thermo-Fisher Scientific). Reverse transcription and cDNA synthesis were carried out by the Maxima First Strand cDNA Synthesis Kit (Thermo-Fisher Scientific). Quantification was performed with the SYBR Select Master Mix (Applied Biosystems) on a CFX96 Touch Real-Time PCR Detection System (Bio-Rad). The small ribosomal subunit protein gene *rpsL* was used as an internal control.

### Antibody production and Western blot

Purified AlgK protein was used as the antigen to generate rabbit polyclonal antiserum using a standard immunization protocol. The anti-AlgK antiserum was further purified by Protein A affinity chromatography (performed by RHBiolabs). A monoclonal rabbit antibody against the bacterial RNA polymerase β-subunit was purchased from Abcam.

An overnight bacterial culture was diluted in 3 mL of alginate-producing medium to an OD_600nm_ of 0.04 and incubated at 37°C for 24 h with agitation at 200 rpm. Cells were harvested by centrifugation and normalized to a final OD_600nm_ of 1.0 in 1 mL PBS buffer. The cell pellet from the normalized resuspension was collected by centrifugation and resuspended in 100 μL of lysis buffer (PBS buffer, 0.1% (w/v) SDS, and 1% (w/v) Triton X-100). Three freeze-thaw cycles using liquid nitrogen were applied to lyse bacteria thoroughly. Cell lysate was then centrifuged at 15,000 g for 10 min, and the supernatant was combined with SDS loading dye. Samples were boiled at 95°C for 10 min prior to loading the clarified lysate onto a 10% (v/v) Tris-Glycine polyacrylamide gel. Following SDS-PAGE, proteins were transferred to a PVDF membrane for immunoblotting. The membrane was blocked using a blocking buffer (5% (w/v) skim milk dissolved in TBST buffer) at 4°C overnight. Blots were washed twice with TBST buffer and the membrane was cut along the 100 kDa molecular weight marker. The membrane containing proteins less than 100 kDa was incubated with anti-AlgK antibody (produced by RHBiolabs) at a 1:2000 dilution in the blocking buffer, while the membrane containing proteins larger than 100 kDa was incubated with anti-RNA polymerase β-subunit antibody (Abcam) at a 1:5000 dilution in the blocking buffer. Both membranes were incubated with primary antibodies at 25°C for 2 h and then washed three times with TBST. The primary antibodies were probed with a goat anti-rabbit horseradish peroxidase (HRP)-conjugated secondary antibody (Abcam) at 1:5000 dilution in the blocking buffer for 2 h at 25°C. Blots were washed three times with TBST. AlgK and RNA polymerase β-subunit bands were detected using the SuperPico ECL Chemiluminescence Kit (Vazyme). Blots were imaged using the ChemiDoc Touch Image System (Bio-Rad).

## Statistical analysis

All raw image data were processed with ImageJ software. Normally distributed data were presented as mean ± SD. Comparisons among three or more groups were performed using one-way ANOVA followed by Tukey’s post hoc test. A *p*-value of less than 0.05 was considered statistically significant. The statistical parameters were shown in the figures and described in the figure legends. All statistical analyses were performed using GraphPad Prism 8.0 (GraphPad Software, USA).

## Data availability

All data described are located within the manuscript and the supporting information. The corresponding atomic models have been deposited to the Electron Microscopy Data Bank (EMDB) and the Protein Data Bank (PDB) under the accession nos. and PDB ID codes: EMD-68393, 22KG (MucD_24-mer); EMD-68645, 22SK (local refinement of trimers from 24-mer); EMD-69903, 24XX (MucD_12-mer); EMD-69907, 24YE (local refinement of trimers from 12-mer).

## Acknowledgements

We thank Prof. Weihui Wu from the College of Life Sciences, Nankai University for the technical assistance.

## Author contributions

Conceptualization: Yujun Jiang, Yong-Gui Gao

Data curation: Yujun Jiang, Xin-Fu Yan

Formal analysis: Yujun Jiang, Xin-Fu Yan

Funding acquisition: Yong-Gui Gao

Investigation: Yujun Jiang, Xin-Fu Yan

Methodology: Yujun Jiang, Chao Wang, Yong-Gui Gao

Software: Yujun Jiang, Xin-Fu Yan

Supervision: Kanaga Sabapathy, Yong-Gui Gao

Validation: Yujun Jiang, Chao Wang, Yong-Gui Gao

Visualization: Yujun Jiang, Xin-Fu Yan, Rya Ero, Chao Wang, Yong-Gui Gao

Writing – original draft: Yujun Jiang, Rya Ero, Yong-Gui Gao

Writing – review & editing: Yujun Jiang, Xin-Fu Yan, Rya Ero, Chao Wang, Yong-Gui Gao

## Supporting Information

**S1 Fig.**
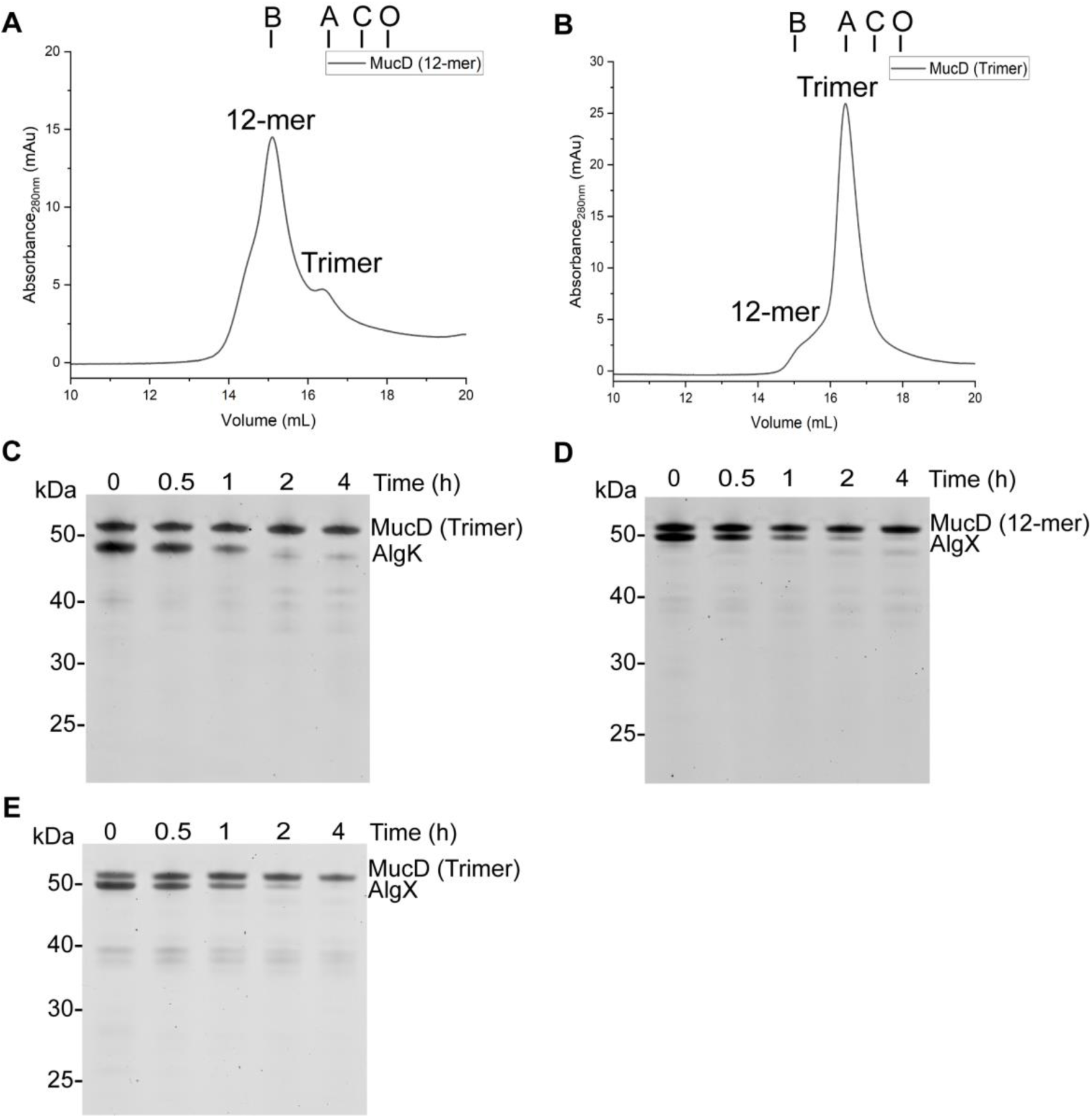
MucD oligomeric state identification and degradation assays. (A and. **B)** Analytical SEC profiles of the MucD 12-mer **(A)** and trimer **(B)**. MucD 12-mer and trimer fractions were isolated from the initial SEC and analyzed separately using a Superose 6 Increase 10/300 GL gel filtration column pre-equilibrated with Buffer E. The theoretical MW of MucD 12-mer is 636 kDa and the trimer is 159 kDa. Elution volumes of molecular weight standards are indicated at the top by vertical lines: bovine thyroglobulin (B), aldolase (A), conalbumin (C), and ovalbumin (O) with molecular weights of 669, 158, 75, and 44 kDa, respectively. **(C, D, and E)** Degradation of AlgK and AlgX by MucD. MucD 12-mer or trimer was incubated with purified AlgK or AlgX in a 1:1 molar ratio at 37°C. At the indicated time points, the aliquots were taken and analyzed by SDS-PAGE.

**S2 Fig.**
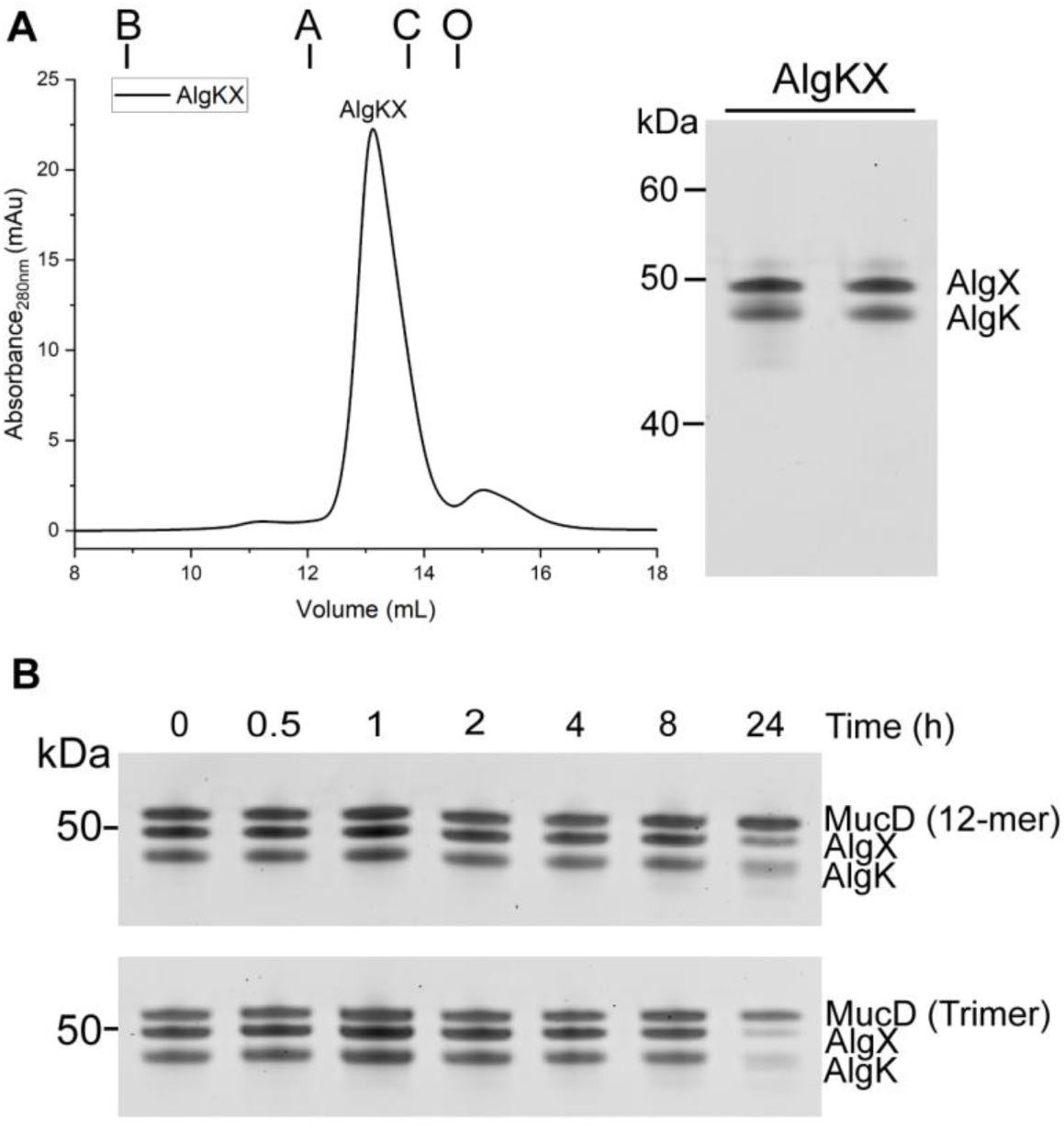
The AlgKX complex formation and degradation assays. **(A)** The formation of AlgKX complex in solution. AlgK and AlgX were incubated in 1:1 molar ratio and the formed complex was purified using SEC with a Superdex 200 Increase 10/300 GL gel filtration column equilibrated with Buffer E (left panel). The complex formation was determined by the calculated molecular weights (MW) based on the SEC calibration curve generated from the gel filtration standard proteins. Fractions corresponding to the AlgKX complex peak were analyzed by SDS-PAGE (right panel). The theoretical MW of AlgK, AlgX, and AlgKX heterodimer are 52.5, 52.6, and 105.1 kDa, respectively. Elution volumes of molecular weight standards are indicated at the top by vertical lines: bovine thyroglobulin (B), aldolase (A), conalbumin (C), and ovalbumin (O) with MW of 669, 158, 75, and 44 kDa, respectively. **(B)** Degradation assays of the AlgKX complex by MucD 12-mer (upper panel) and trimer (lower panel). MucD 12-mer or trimer was incubated with the AlgKX complex in a 1:1 molar ratio at 37°C. At shown time points, the aliquots were taken and analyzed by SDS-PAGE.

**S3 Fig.**
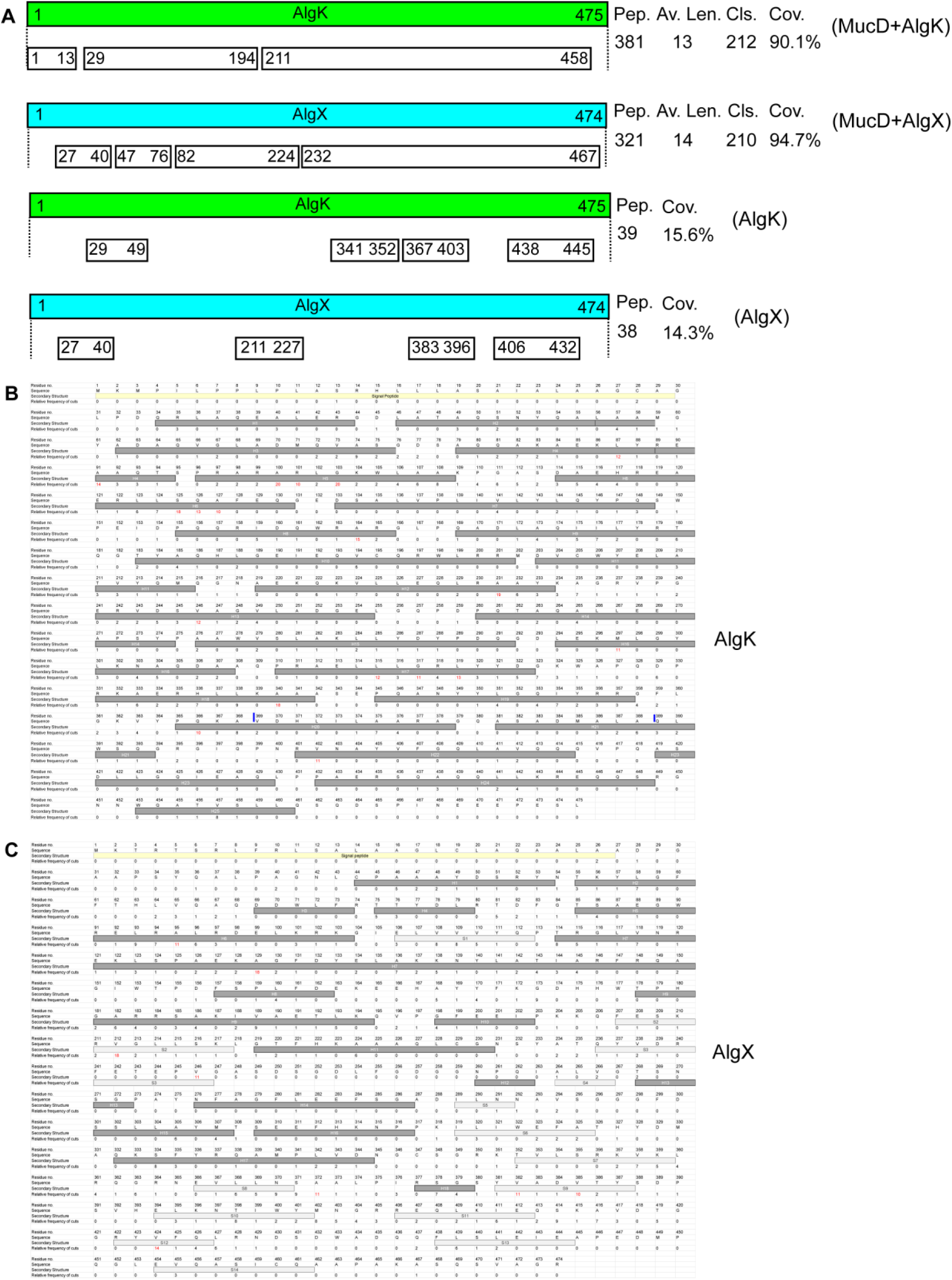
Analysis of mass spectrometry data of AlgK and AlgX degradation products. **(A)** Overview of the degradation products of AlgK and AlgX detected by LC-MS. Green bars: linear representation of AlgK primary structure. Blue bars: linear representation of AlgX primary structure. Detected peptides from the incubation of MucD with AlgK (MucD+AlgK), MucD with AlgX (MucD+AlgX), AlgK only (AlgK) or AlgX only (AlgX), whose sequences aligned without gaps were grouped into fragments and shown as white bars below. The length of bars is disproportionate for clearer illustration. Pep.: the total number of detected peptides. Av. Len.: the average length of all detected peptides. Cls.: the total number of cleavage sites. Cov.: the coverage of the AlgK or AlgX sequence by detected peptides. **(B and C)** The relative frequency of cuts (RFC) at each amino acid residue was calculated and aligned with AlgK **(B)** and AlgX **(C)** primary sequences. Secondary structure elements such as helices (H), sheets (S), and loops (left as blank) were labelled based on the AlphaFold predicted AlgK model (ID: AF-P96956-F1) and the solved AlgX structure (PDB: 4KNC). Amino acid residues with RFC > 9 were considered as high RFC sites and were highlighted in red. RFC=0 indicated no detected cleavage. The sequence of the synthesized polypeptide (AlgK_369-388_) for structural determination was highlighted by two blue lines. The data represents average results from three independent experiments.

**S4 Fig.**
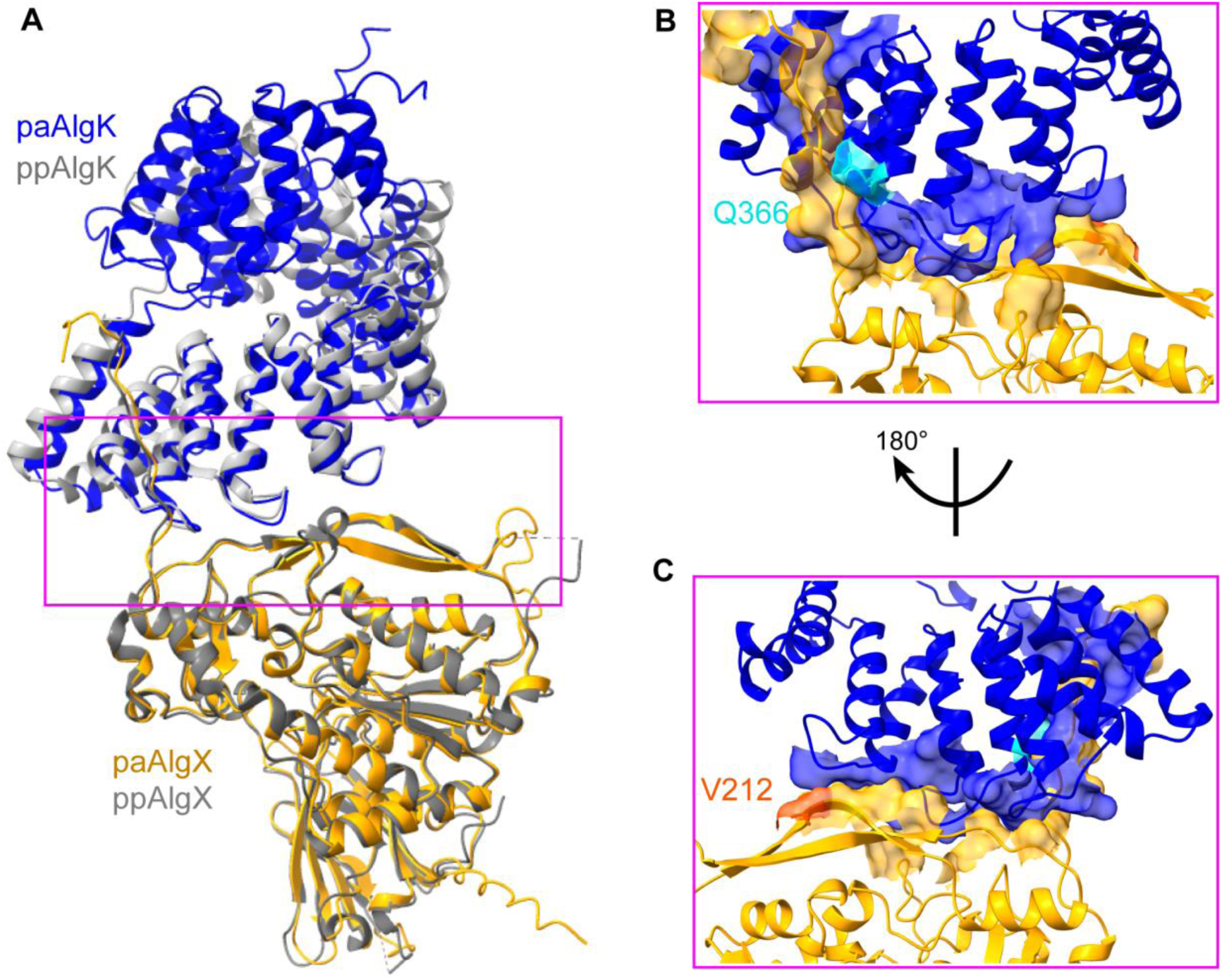
Structural alignment of high RFC residues at the AlgKX complex interface. **(A)** Structural comparison of AlphaFold predicted paAlgKX (from *P. aeruginosa*) complex with ppAlgKX (from *P. putida*, PDB: 7ULA). Two models are superimposed together, with Cα RMSDs of 0.68 Å. The paAlgK model is shown in blue and paAlgX in orange. The pink rectangle highlights the AlgKX interface. **(B and C)** Close-up view of the paAlgKX complex interface overlaid with the high RFC residues. Residues involved in the paAlgKX complex interactions are shown in surface as well, while Q366 (RFC=10) from paAlgK is colored in cyan and V212 (RFC=18) from paAlgX is colored in orange red.

**S5 Fig.**
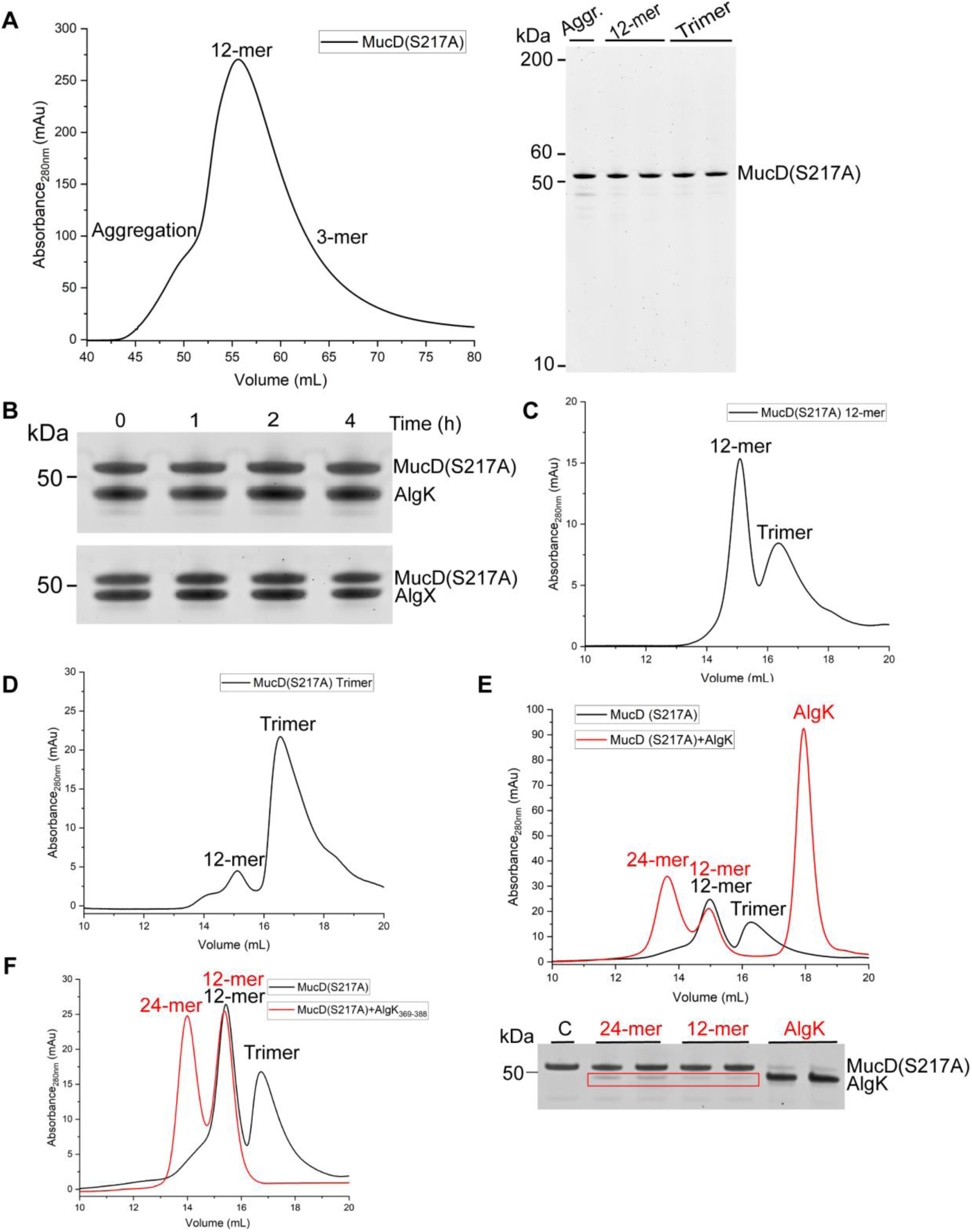
MucD(S217A) purification, analysis and complex formation with the substrates. **(A)** The SEC profile of MucD(S217A). MucD(S217A) was loaded onto a HiLoad 16/600 Superdex 200 gel filtration column equilibrated with Buffer E. Fractions corresponding to peaks (left panel) were analyzed by SDS-PAGE (right panel). **(B)** AlgK and AlgX degradation assays by MucD(S217A) 12-mer. MucD(S217A) was incubated with AlgK or AlgX in a 1:1 molar ratio at 37°C. At indicated time points, the aliquots were taken and analyzed by SDS-PAGE. **(C and D)** Analytical SEC profiles of MucD(S217A) 12-mer **(C)** and trimer **(D)**. MucD(S217A) 12-mer and trimer fractions were isolated from the initial SEC and analyzed separately using a Superose 6 Increase 10/300 GL gel filtration column pre-equilibrated with Buffer E. **(E)** MucD(S217A) complex formation with AlgK. MucD(S217A) 12-mer was incubated with AlgK at a 1:1 molar ratio (120 μM) for 2 h at 37°C. The sample was then analyzed using a Superose 6 Increase 10/300 GL gel filtration column pre-equilibrated with Buffer S. Peak fractions were analyzed by SDS-PAGE. Co-eluted AlgK bands with MucD are highlighted by a red rectangle. The lane “C”: MucD(S217A) 12-mer without AlgK addition serves as the control. **(F)** MucD(S217A) complex formation with AlgK_369-388_. MucD(S217A) 12-mer was incubated with AlgK_369-388_ as described in Methods. The sample was then analyzed using a Superose 6 Increase 10/300 GL gel filtration column pre-equilibrated with Buffer S. The oligomeric states were determined by the calculated molecular weights (MW) based on the SEC calibration curve generated from the gel filtration standard proteins. The theoretical MW for MucD(S217A) 24-mer, MucD(S217A) 12-mer, and MucD(S217A) trimer are 1,272 kDa, 636 kDa and 159 kDa, respectively.

**S6 Fig.**
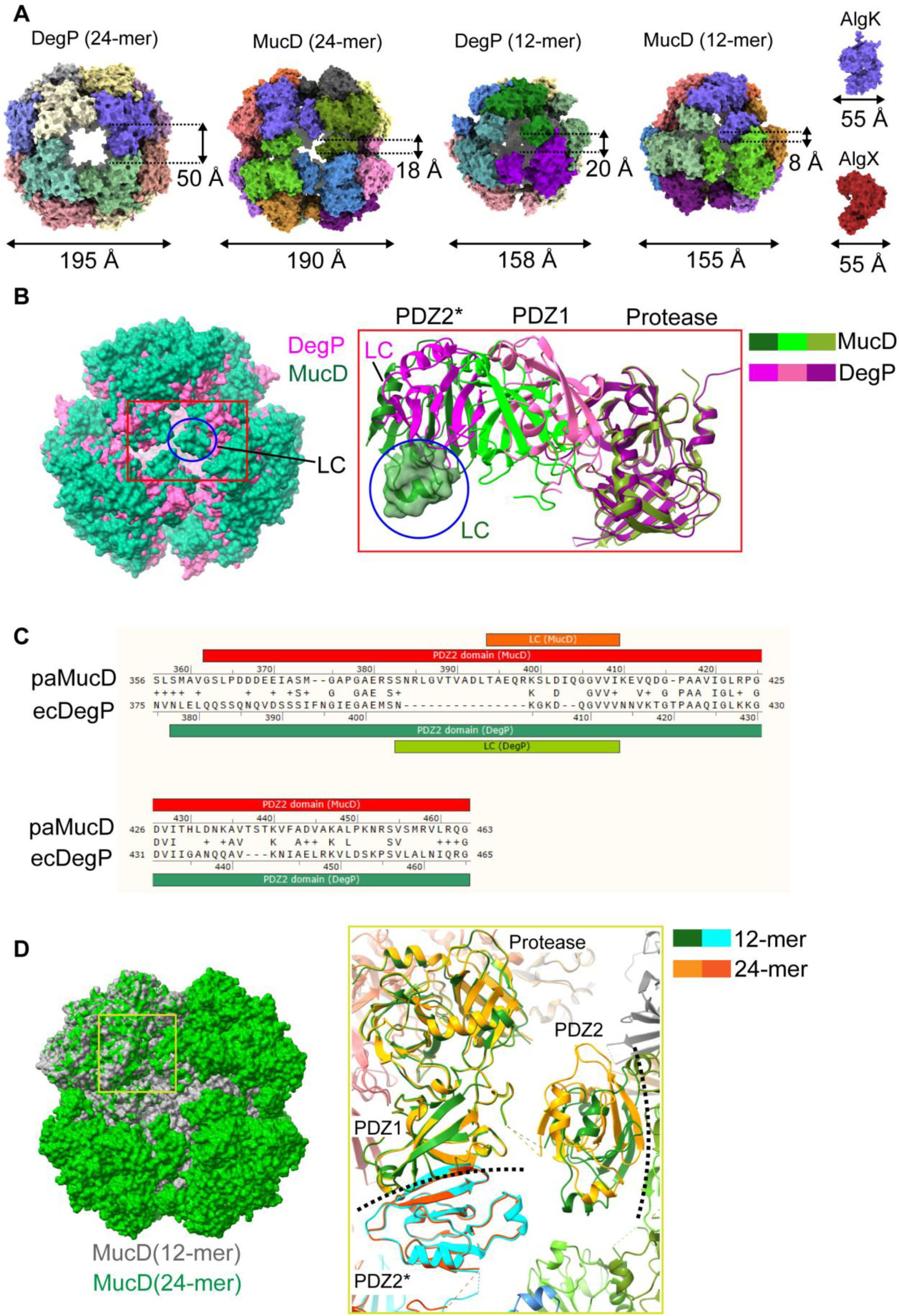
Comparison of MucD and DegP higher-order oligomers assembly. **(A)** Higher-order oligomers assembly comparison of DegP 24-mer (PDB: 3OU0), MucD 24-mer (PDB: 22KG), DegP 12-mer (PDB: 8F0U), and MucD 12-mer (PDB: 24XX). AlgK (ID: AF-P96956-F1) and AlgX (PDB: 4KNC) models are shown together for size references. The surface representations of all models are at the same scale. Each protomer of MucD and DegP cages is colored differently. **(B)** Structural alignment of 12-mer DegP and 12-mer MucD. The two models are shown as surfaces, and domains in close proximity to the surface pores are highlighted with a cyan rectangle, with close-up views shown in cartoons (right panel), respectively. The “cage loop” (LC) of MucD is highlighted by a blue oval (right panel) and also shown in surface representation in the close-up view. PDZ2*: the PDZ2 domain is from a neighbouring trimer. **(C)** Sequence alignment of MucD and DegP PDZ2 domains by Snapgene software. The LC length of paMucD (from *P. aeruginosa*) and ecDegP (from *E. coli*) are 16 and 10 amino acid residues, respectively. +: Similar residues. Blank: Not similar. **(D)** Structural alignment of MucD 12-mer and 24-mer. The two models are shown as surfaces in green (24-mer) and grey (12-mer) (left panel). A close-up view of protomers of two trimers at the inter-trimer interface is shown in cartoons (right panel). The flexible linker between the consecutive PDZ1 and PDZ2 domains is indicated by colored dotted lines. All domains within the same trimer are colored the same. The black dotted lines indicate the inter-trimer interfaces.

**S7 Fig.**
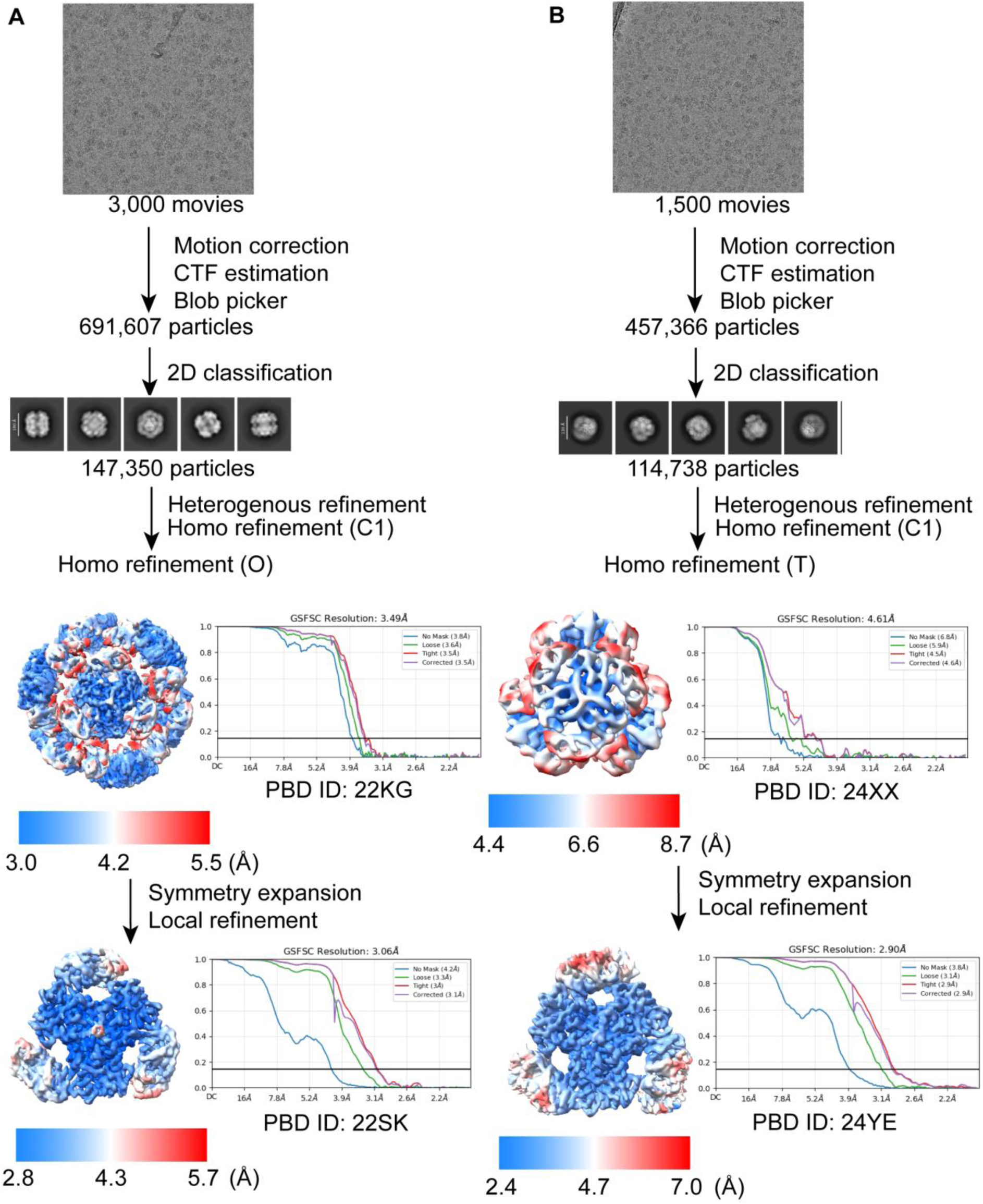
Cryo-EM data processing of MucD(S217A)-AlgK_369-388_. (A and. **B)** The workflow for the 24-mer **(A)** and 12-mer **(B)** of MucD(S217A)-AlgK_369-388_ complex dataset processing and the local resolution estimation of electron density maps. The representative micrographs, 2D classes and final FSC curves are shown as well.

**S8 Fig.**
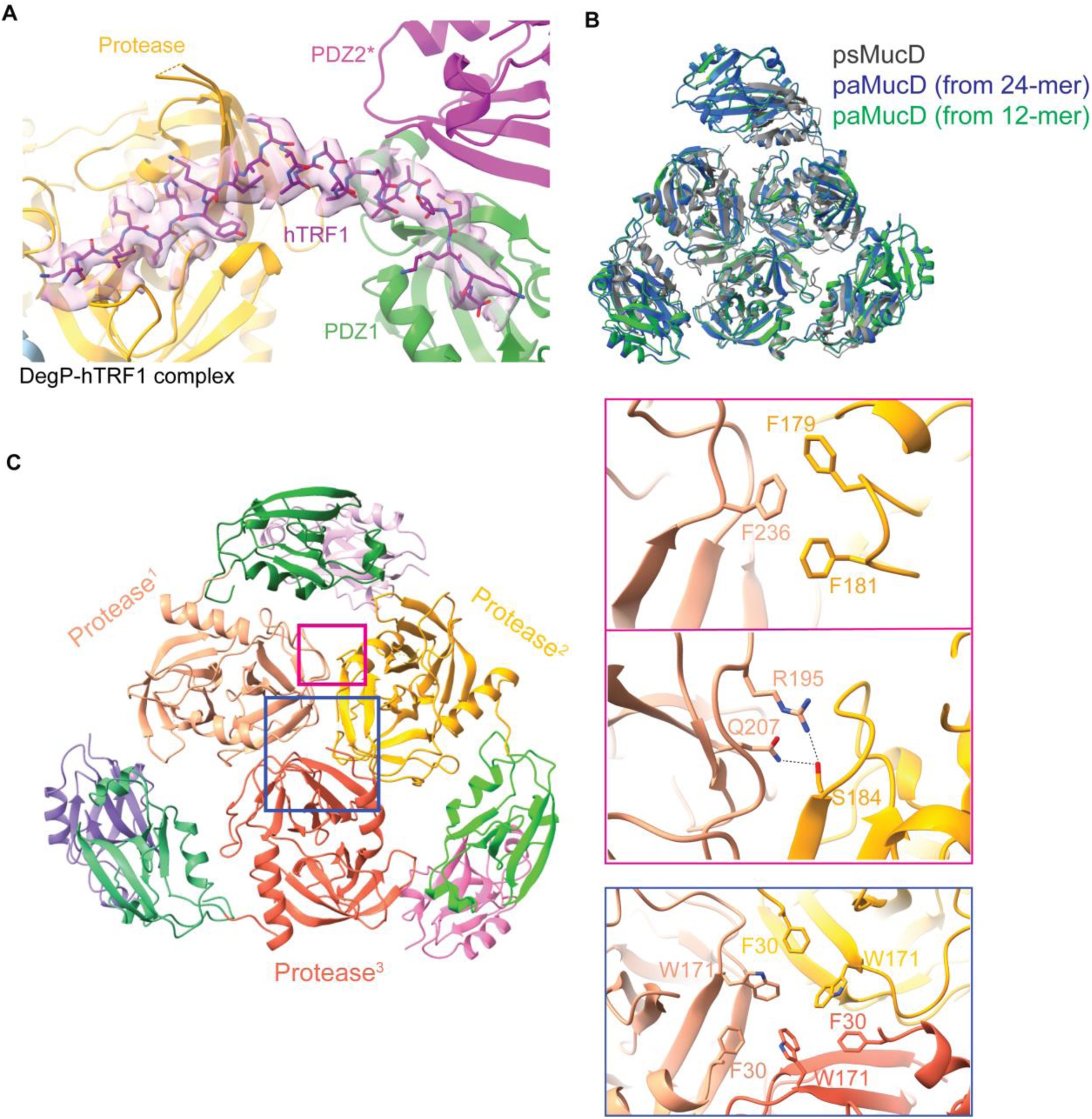
Structural analysis of the trimeric MucD. **(A)** Model representation of DegP-hTRF1 complex (PDB: 8F0A) structure. The bound substrate hTRF1 with DegP is shown in sticks and fitted into the electron density map. **(B)** Structural comparison of trimeric states of psMucD (from *P. syringae*, PDB: 8K2Y) and paMucD (the locally refined trimers of 24-mer from *P. aeruginosa*, PDB: 22SK; the locally refined trimers of 12-mer from *P. aeruginosa*, PDB: 24YE). The Cα RMSD value of psMucD and paMucD (from 24-mer) is 0.69 Å while the value of paMucD (from 24-mer) and paMucD (from 12-mer) is 0.49 Å. **(C)** Characterization of MucD intra-trimer interactions. The protease domains that are involved in intra-trimer interactions are labelled. Blue rectangle: Close-up view of interacting residues F30 and W171 (shown in sticks) that are involved in joining the three protomers. Pink rectangles: close-up views of the residues F179-F181 and F236 (upper box) and R195, Q207, and S184 (lower box) that interact at the interface of two nearby protomers.

**S9 Fig.**
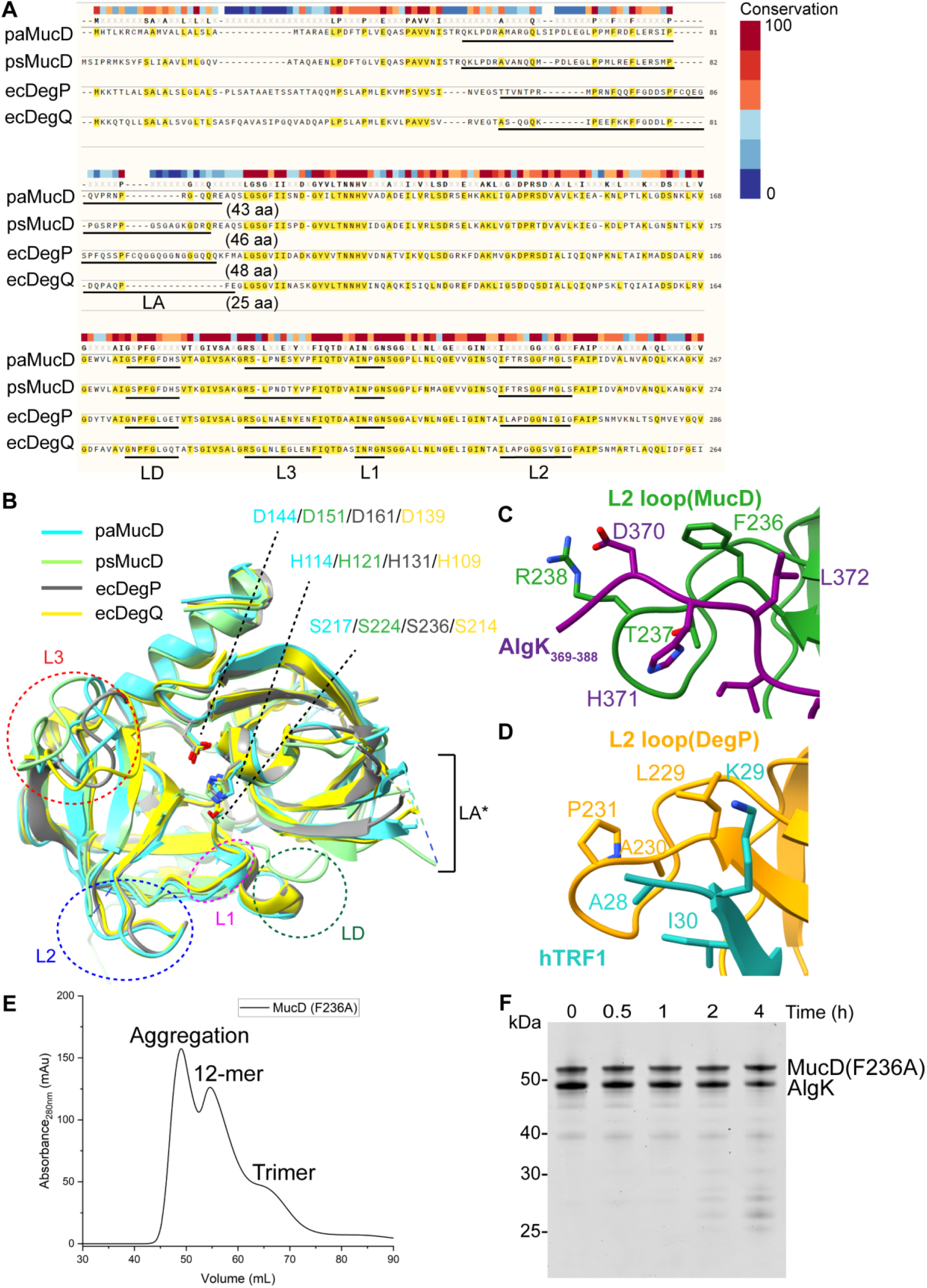
Sequence and structural alignment of MucD protease domains with other bacterial HtrA-like proteases. **(A)** Sequence alignment of paMucD (from *P. aeruginosa*), psMucD (from *P. syringae*), ecDegP and ecDegQ (from *E. coli*). The amino acids of LA, LD, L1, L2, and L3 loops are underlined, and the length of the LA loops is labelled aside. The alignment and conservation score calculation was conducted by SnapGene software. Amino acids that match the reference (paMucD) are marked with yellow highlighting. **(B)** Structural alignment of the protease domain of paMucD (PDB: 22SK), psMucD (PDB: 8K2Y), ecDegP (PDB: 3OU0), and ecDegQ (PDB: 3STJ). The conserved L1, L2, L3, and LD loops are highlighted with dotted ovals. LA*: The LA loops of all four structures were unable to be modelled due to their flexibility. The amino acid residues at the two ends of the missing structures are connected by dotted lines. The catalytic triad residues are shown as sticks and labelled. **(C and D)** Comparison of interaction between bound substrates and the L2 loop of MucD **(C)** (PDB: 22SK) or DegP **(D)** (PDB: 8F0A). The interacting residues are shown as sticks and labelled. **(E)** The SEC profile of MucD(F236A). The purified protein was loaded onto a HiLoad 16/600 Superdex 200 gel filtration column, and the absorbance of elution was recorded at 280 nm. **(F)** Degradation of AlgK by MucD(F236A). The MucD(F236A) was incubated in a 1:1 molar ratio with purified AlgK at 37°C. At the indicated time points, aliquots were taken and analyzed by SDS-PAGE.

**S10 Fig.**
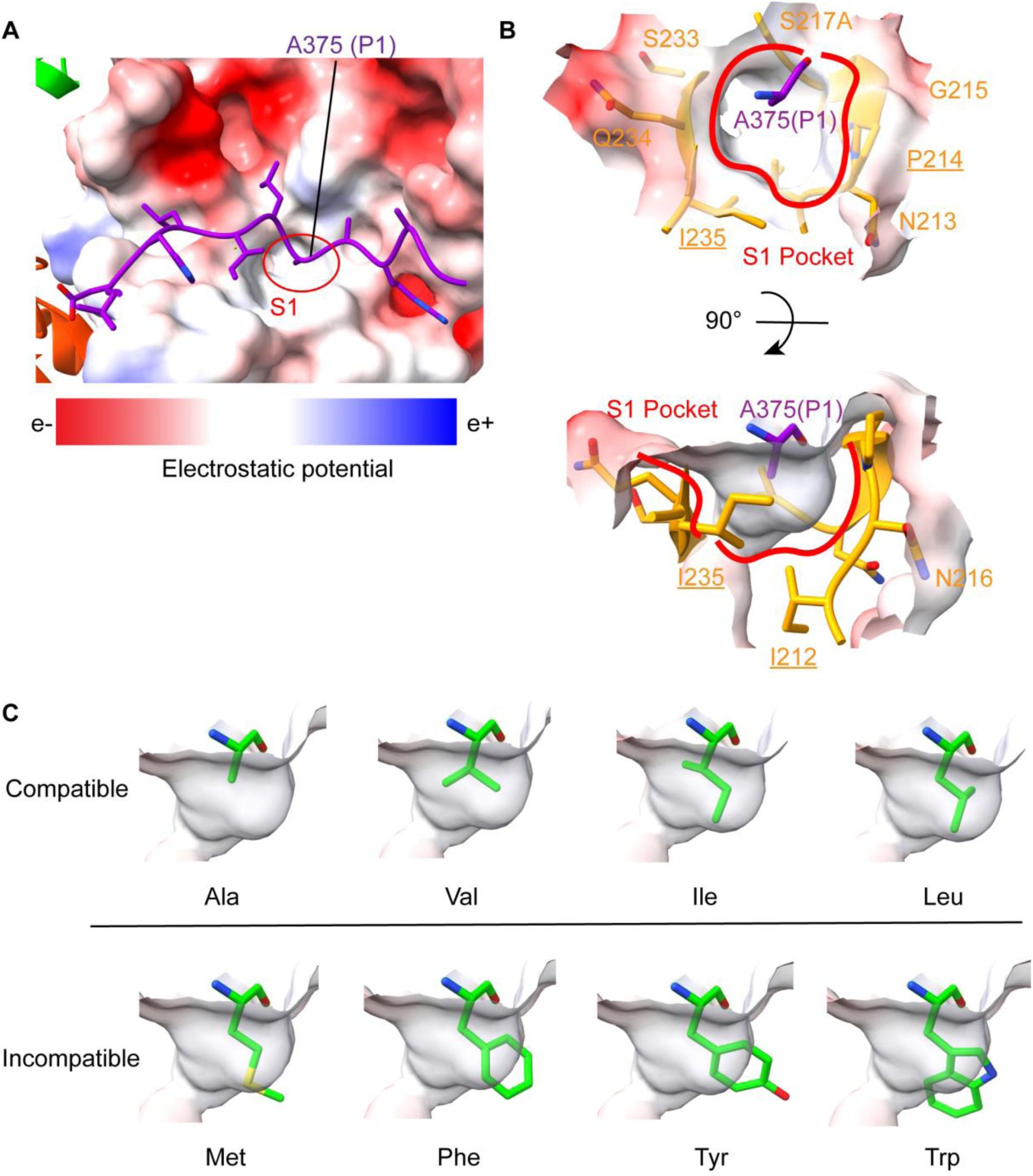
Interaction of the substrate P1 residue with the MucD S1 pocket. **(A)** Electrostatic surface representation of the MucD protease domain. The bound peptide AlgK_369-388_ is shown in purple sticks. The corresponding hydrophobic S1 pocket is highlighted by a red oval. The electrostatic potential scale ranges from-11.3 kT/e (red) to +11.3 kT/e (blue). **(B)** Close-up views of the P1-S1 interaction. The S1 pocket is shown in electrostatic surface and highlighted by a red ring while the P1 residue (A375 of AlgK_369-388_) is shown as purple sticks. The residues of the MucD protease domain that involved in the S1 pocket formation are shown in orange sticks. Key residues forming the S1 pocket are underlined. **(C)** Compatibility of hydrophobic amino acids with the S1 pocket of MucD. The S1 pockets are shown in electrostatic surface. The P1 residue (A375 of AlgK_369-388_) is substituted with other hydrophobic rotamer residues, which are shown in green sticks. Compatible: the side chains of the P1 residue fit in the S1 pocket. Incompatible: the side chains of the P1 residue exceed the size limit of the S1 pocket and cause spatial clash.

**S11 Fig.**
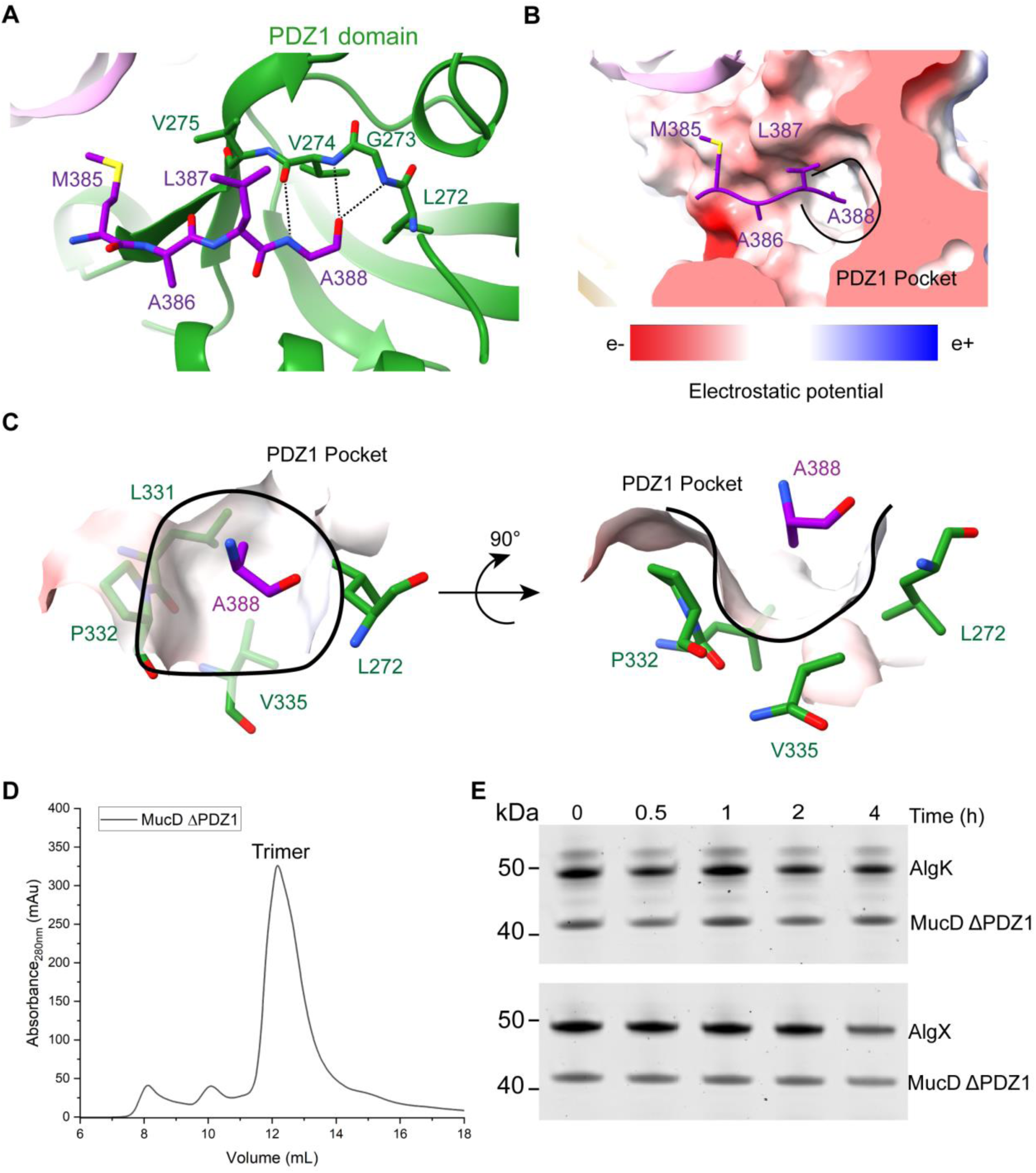
Interaction of the MucD PDZ1 domain with the bound AlgK_369-388_ peptide. **(A)** Cartoon representation of the AlgK_369-388_-PDZ1 interaction. Four residues from the C-terminus of AlgK_369-388_ that had clear electron density are labelled and shown in purple sticks, while the PDZ1 residues involved in the interaction are displayed as green sticks. Dotted black lines represent the backbone hydrogen bonds. **(B)** Electrostatic surface representation of the MucD PDZ1 domain. The corresponding PDZ1 pocket is highlighted by a black ring and the substrate residues are shown in purple sticks. The electrostatic potential scale ranges from-11.3 kT/e (red) to +11.3 kT/e (blue). **(C)** A close-up view of the A388-PDZ1 pocket interaction. The PDZ1 pocket is shown in electrostatic surface and highlighted by the black ring. The residues of the MucD PDZ1 domain that are involved in the pocket formation are shown in green sticks while the C-terminal residue A388 of AlgK_369-388_ is shown in purple sticks. **(D)** The SEC profile of purified MucD ΔPDZ1. Purified MucD ΔPDZ1 was loaded onto a Superdex 200 Increase 10/300 GL gel filtration column pre-equilibrated with Buffer E. The oligomeric state was determined by the calculated molecular weights (MW) based on the SEC calibration curve generated from the gel filtration using standard proteins. The theoretical MW of MucD ΔPDZ1 is 41 kDa. **(E)** Degradation assays of AlgK and AlgX by MucD ΔPDZ1. Degradation assays were conducted as described in Methods. At indicated time points, an aliquot was taken and analyzed by SDS-PAGE.

**S12 Fig.**
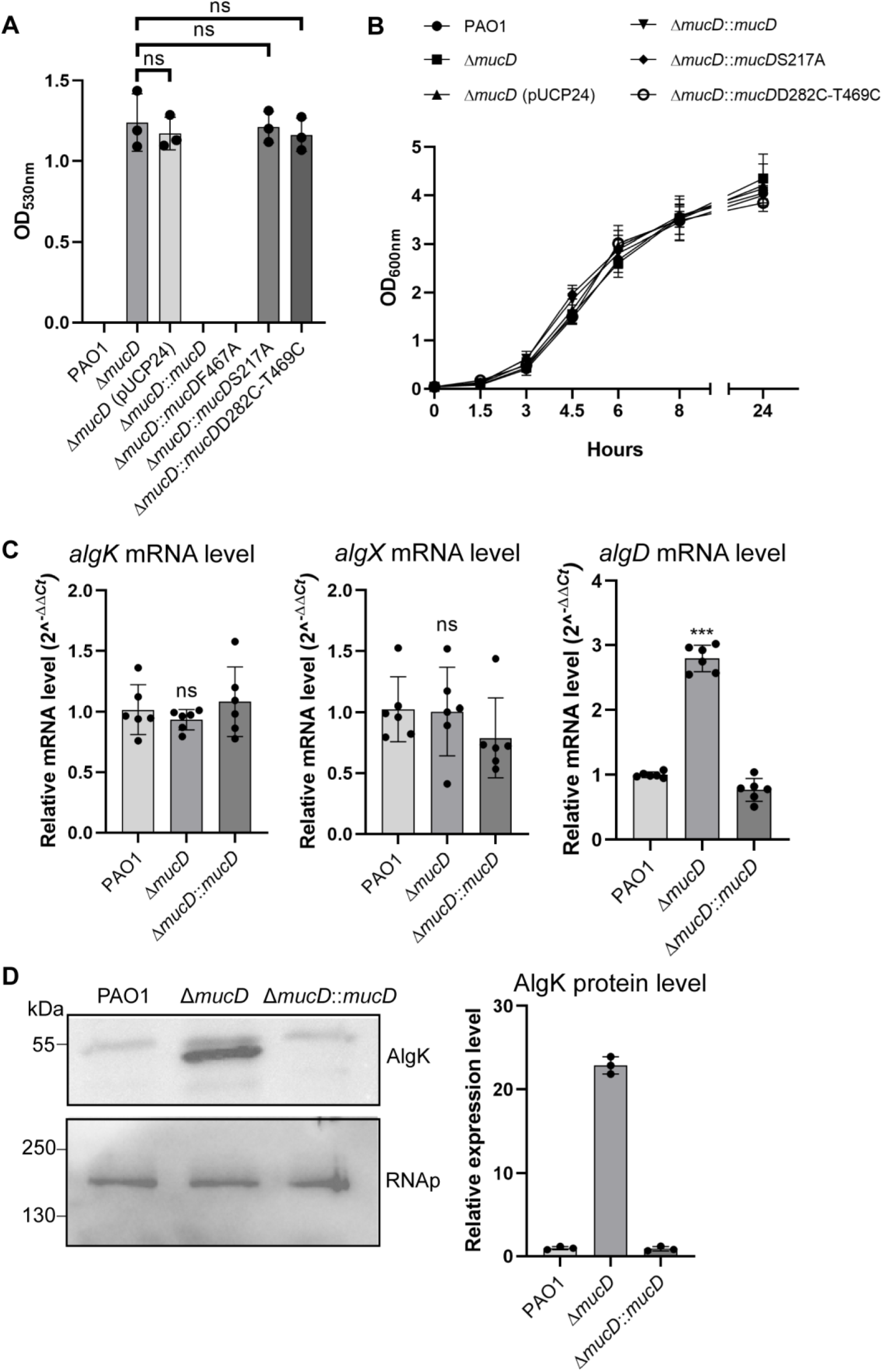
Regulation of alginate production by MucD in *P. aeruginosa*. **(A)** Alginate production detection in *P. aeruginosa* PAO1 strain variants. The chart combines results from three independent experiments with error bars representing the standard deviations. Statistically significant variations calculated using the one-way ANOVA test are shown above the corresponding bars. ns: not significant (P>0.05) **(B)** Growth curves of *mucD*-related mutants in the PAO1 background. The cell density was measured by OD_600nm_ at indicated time points. The data represent results from three independent experiments and error bars represent the standard deviations. **(C)** The *algK*, *algX,* and *algD* mRNA levels in PAO1, *ΔmucD,* and *ΔmucD*::*mucD* strains. Total RNA was isolated and the mRNA levels of all genes were determined using quantitative reverse transcription PCR (RT-qPCR). The fold change of transcriptional level was normalized to PAO1. All shown results were from three independent experiments with error bars representing the standard deviations. Statistically significant variations calculated using the one-way ANOVA test are shown above the corresponding bars. ns: not significant (P>0.05); ***: P<0.001. **(D)** AlgK protein expression levels in PAO1, *ΔmucD,* and *ΔmucD*::*mucD* strains. Whole cell lysis was used for the AlgK protein detection by Western blot with an anti-AlgK polyclonal antibody. RNA polymerase β-subunit (RNAp) served as a loading control. Detection of AlgK and RNAp was performed on the same blot. The intensity of AlgK protein bands was quantified using ImageJ software and the fold change of expression level was normalized to PAO1. A representative result from three independent experiments is shown.

**S13 Fig.**
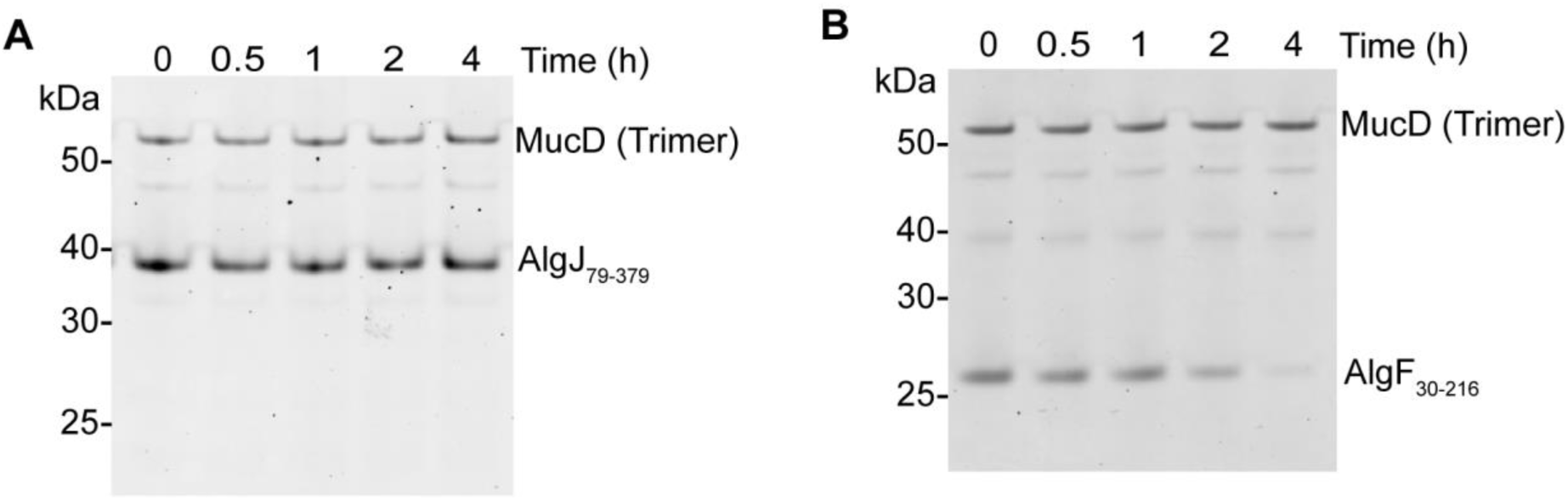
Degradation assays of AlgJ_79-379_ and AlgF_30-216_ by MucD. MucD trimer was incubated with purified AlgJ_79-379_ **(A)** or AlgF_30-216_ **(B)** at 37°C in a 1:1 molar ratio. At the indicated time points, the aliquots were taken and analyzed by SDS-PAGE.

**S14 Fig.**
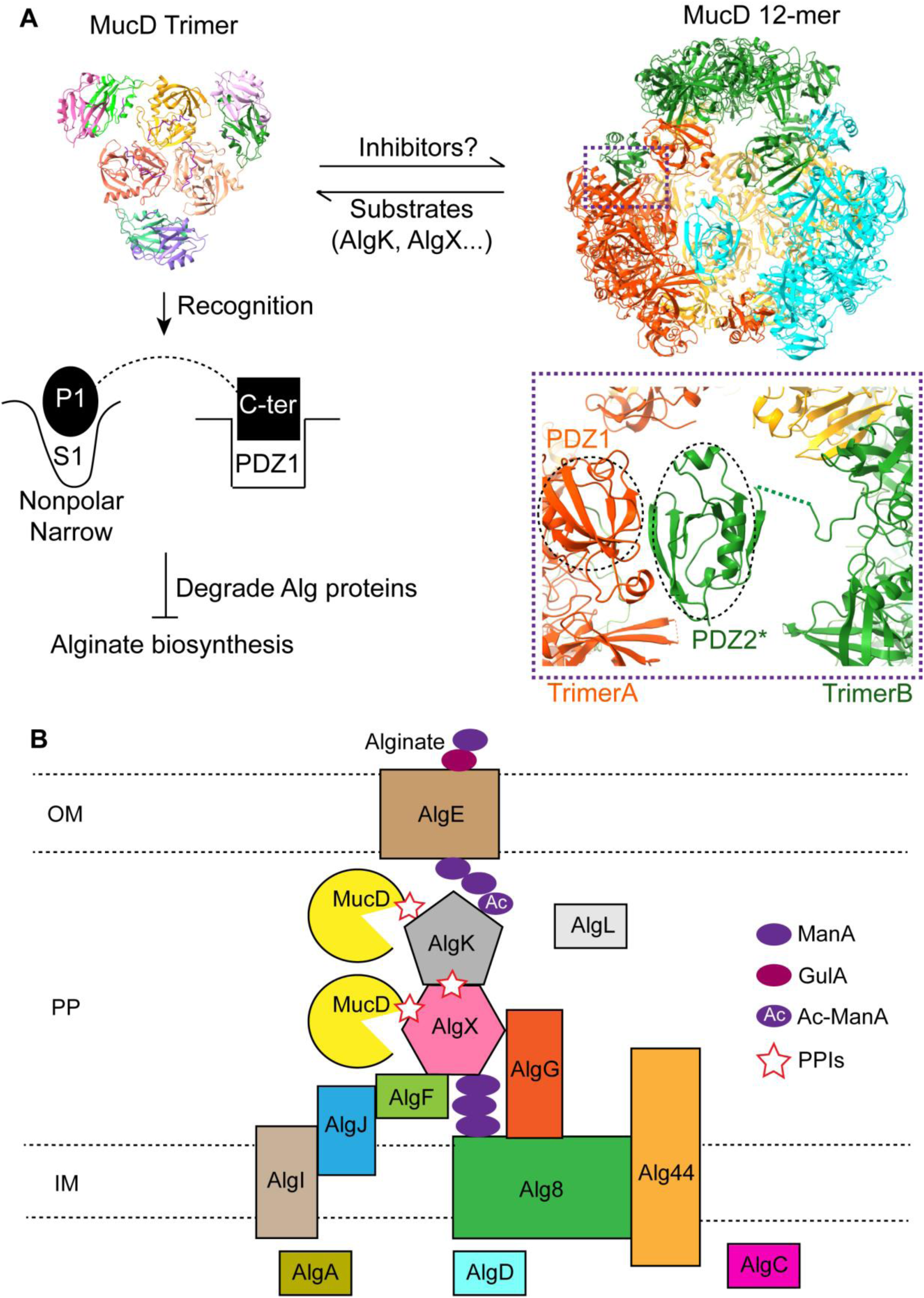
Model representation of MucD regulation in alginate biosynthesis. **(B)** A schematic model for MucD-mediated regulation of alginate biosynthesis. *In vivo*, MucD is proposed to equilibrate between 12-mer and trimeric states. Upon substrate proteins (i.e., AlgK and AlgX) presence, the active trimer state carries out proteolysis and causes the equilibrium to shift from the 12-mer to the trimer, leading to the negative regulation of alginate biosynthesis. After substrate degradation is completed, potential inhibitors (such as serpins and small protein inhibitors) may bind to trimers and restore the resting 12-mer state via PDZ1-PDZ2* interaction. PDZ2*: the PDZ2 domain is from a neighbouring trimer. **(B)** A schematic representation of the proposed interaction network within the alginate biosynthesis complex. Proteins positioned adjacent to one another are proposed to form interactions. Star symbols highlight the interactions among MucD, AlgK, and AlgX, emphasizing the relevance of the present findings to MucD-mediated regulation of alginate biosynthesis. IM, inner membrane; PP, periplasm; OM, outer membrane; ManA, β-D-mannuronate; GulA, α-L-guluronate; Ac-ManA, O-acetylated β-D-mannuronate; PPIs, protein-protein interactions.

---

**S1 Table.**
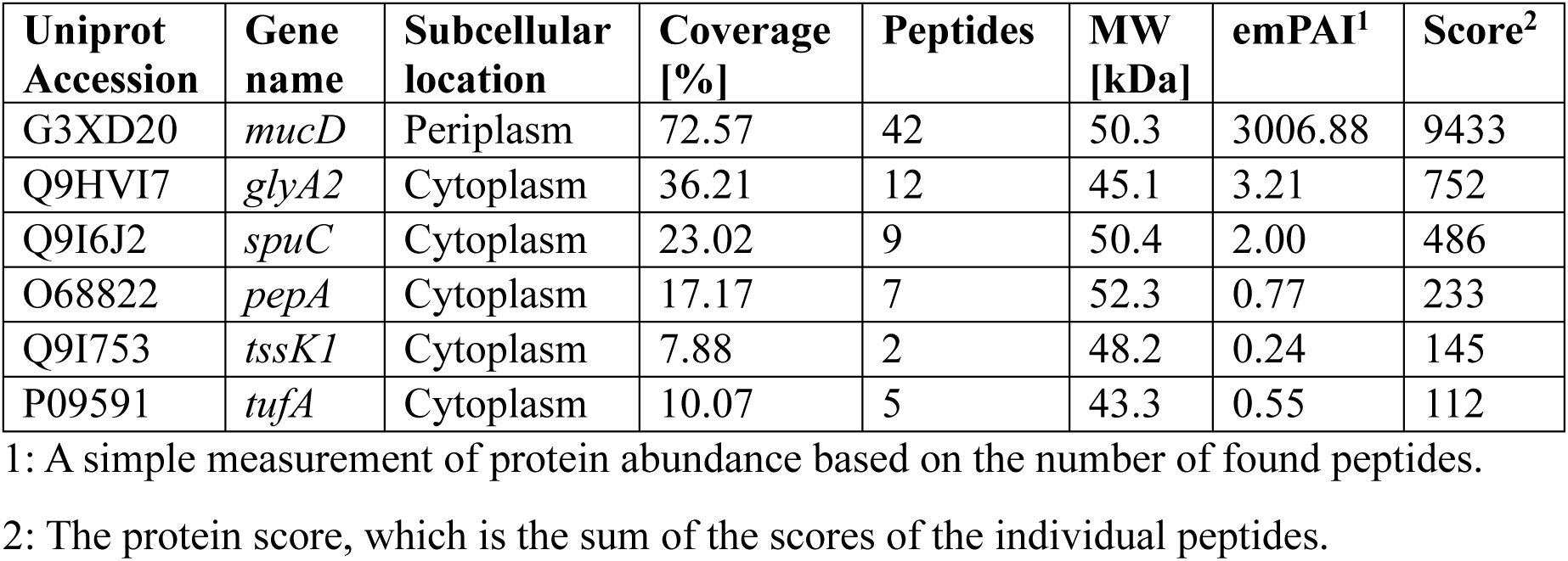
MALDI-TOF/MS identification of purified MucD.

**S2 Table.** Cryo-EM data collection, refinement, and validation statistics.

|  | **MucD 24-mer & trimers**<br>**local refinement with the**<br>**client peptide** |  | **MucD 12-mer & trimers**<br>**local refinement with the**<br>**client peptide** |  |
| --- | --- | --- | --- | --- |
| PDB entry ID | 22KG | 22SK | 24XX | 24YE |
| **Data collection** |  |  |  |  |
| Magnification | 130,000 | 130,000 | 130,000 | 130,000 |
| Pixel size (Å) | 0.97 | 0.97 | 0.97 | 0.97 |
| Defocus range (µm) | -0.6 ~ -1.2 | -0.6 ~ -1.2 | -0.6 ~ -1.2 | -0.6 ~ -1.2 |
| Voltage (kV) | 300 | 300 | 300 | 300 |
| Electron dose (e<sup>-</sup> /Å<sup>2</sup>) | 40 | 40 | 40 | 40 |
| Particles | 147,350 | 147,350 | 114,738 | 114,738 |
| **Refinement** |  |  |  |  |
| Average Resolution (Å) | 3.49 | 3.06 | 4.61 | 2.90 |
| Map sharpening B-factor (Å<sup>2</sup>) | 140.0 | 68.6 | 194.7 | 65.2 |
| **Model composition** |  |  |  |  |
| Non-hydrogen atoms | 70,032 | 9,069 | 35,016 | 9,048 |
| Protein residues | 9432 | 1,224 | 4,716 | 1,221 |
| **ADP (B-factors) (Å<sup>2</sup>)** |  |  |  |  |
| Protein (min/max/mean) | 60.13/169.48/<br>111.18 | 30.00/169.48<br>/111.37 | 21.25/237.66<br>/108.87 | 21.22/238.36/<br>107.41 |
| Bond lengths (Å) | 0.007 | 0.008 | 0.004 | 0.005 |
| Bond angles (°) | 1.121 | 1.128 | 1.061 | 1.057 |
| **Validation** |  |  |  |  |
| Molprobity score | 1.79 | 1.95 | 1.87 | 2.04 |
| Clashscore | 9.89 | 13.45 | 12.09 | 16.54 |
| Rotamer outliers (%) | 0.00 | 0.10 | 0.05 | 0.10 |
| **Ramachandran plot (%)** |  |  |  |  |
| Favoured | 96.02 | 95.56 | 96.04 | 95.38 |
| Allowed | 3.98 | 4.36 | 3.96 | 4.62 |
| Outliers | 0.00 | 0.08 | 0.00 | 0.00 |

**S3 Table.** MALDI-TOF/MS identification of partially degraded products of AlgK.

| **Uniport Accession** | **Gene name** | **Subcellular location** | **Coverage [%]** | **Peptides** | **MW [kDa]** | **emPAI<sup>1</sup>** | **Score<sup>2</sup>** |
| --- | --- | --- | --- | --- | --- | --- | --- |
| P96956 | *algK* | Periplasm | 28.63 | 21 | 52.4 | 19.89 | 1972 |

1: A simple measurement of protein abundance that is based on the number of found peptides.
2: The protein score, which is the sum of the scores of the individual peptides.

**S4 Table.** Bacterial strains and plasmids used in this study.

| Strains or plasmids | Description | Source |
| --- | --- | --- |
| **Strains** |  |  |
| DH5α | F<sup>-</sup> $\phi$ 80*lacZ*ΔM15 *endA1 recA1 hsdR17*(rK<sup>-</sup> mK<sup>+</sup>) *supE44 thi-1 relA1* $\Delta$(*lacZYA-argF*)U169 *gyrA96 deoR* | Thermo-Fisher Scientific |
| BL21 (DE3) | *fhuA2 [lon] ompT gal* ($\lambda$ DE3) [*dcm*] Δ*hsdS* $\lambda$ DE3 = $\lambda$ *sBamHI* Δ*EcoRI-B* *int::(lacI::PlacUV5::T7 gene1)* *i21* Δ*nin5* | NEB |
| S17-1 | RP4-2 Tc::Mu Km::Tn7 Tp<sup>r</sup> Sm<sup>r</sup> Pro Res<sup>-</sup> Mod<sup>+</sup> | [1] |
| PAO1 | Wild type *P. aeruginosa* strain | [2] |
| PAO1 Δ*mucD* | PAO1 with *mucD* gene deletion | This study |
| **Plasmids** |  |  |
| pUCP24 | Shuttle vector between *E. coli* and *P. aeruginosa*; Gentamicin<sup>r</sup> | [1] |
| pEX18Tc | Gene knockout vector; Tetracycline<sup>r</sup> | [3] |
| pMMB67EH | IPTG-inducible expression vector for *Pseudomonas*; Carbenicillin<sup>r</sup> | [4] |
| pDBHT-2 | IPTG-inducible expression vector for *E. coli*; Kanamycin<sup>r</sup> | [5] |
| pUCP24-*mucD* | *mucD* gene in pUCP24 | This study |
| pUCP24-*mucD*(S217A) | *mucD*(S217A) gene in pUCP24 | This study |
| pUCP24-*mucD*(F467A) | *mucD*(F467A) gene in pUCP24 | This study |
| pUCP24-*mucD*(D282C-T469C) | *mucD*(D282C-T469C) gene in pUCP24 | This study |
| pMMB67EH-MucD-6×His | 6 x His-tagged MucD protein on pMMB67EH | This study |
| pMMB67EH-MucD(S217A)-6×His | 6 x His-tagged MucD protein with S217A mutation on pMMB67EH | This study |
| pMMB67EH-MucD(F467A)-6×His | 6 x His-tagged MucD protein with F467A mutation on pMMB67EH | This study |
| pMMB67EH-MucD(D282C-T469C)-6×His | 6 x His-tagged MucD protein with D282C-T469C mutations on pMMB67EH | This study |
| pMMB67EH-AlgK-6×His | 6 x His-tagged AlgK protein on pMMB67EH | This study |
| pMMB67EH-AlgX-6×His | 6 x His-tagged AlgX protein on pMMB67EH | This study |
| pDBHT-2-6×His-AlgF<sub>30-216</sub> | 6 x His-tagged AlgF<sub>30-216</sub> protein on pDBHT-2 | This study |
| pDBHT-2-6×His-AlgJ<sub>79-379</sub> | 6 x His-tagged AlgJ<sub>79-379</sub> protein on pDBHT-2 | This study |
| pEX18Tc-*mucD*-dele | Upstream and downstream regions flanking *mucD* gene for gene deletion on pEX18Tc | This study |

## References

1. Hauser AR. The type III secretion system of Pseudomonas aeruginosa: infection by injection. Nat Rev Microbiol. 2009;7: 654–665. doi:10.1038/nrmicro2199

2. Boles BR, Thoendel M, Singh PK. Self-generated diversity produces “insurance effects” in biofilm communities. Proc Natl Acad Sci U S A. 2004;101: 16630–16635. doi:10.1073/pnas.0407460101

3. Whitney JC, Howell PL. Synthase-dependent exopolysaccharide secretion in Gram-negative bacteria. Trends in Microbiology. 2013;21: 63–72. doi:10.1016/j.tim.2012.10.001

4. Boucher JC, Yu H, Mudd MH, Deretic V. Mucoid Pseudomonas aeruginosa in cystic fibrosis: characterization of muc mutations in clinical isolates and analysis of clearance in a mouse model of respiratory infection. Infect Immun. 1997;65: 3838–3846. doi:10.1128/iai.65.9.3838-3846.1997

5. Lyu ZX, Zhao XS. Periplasmic quality control in biogenesis of outer membrane proteins. Biochemical Society Transactions. 2015;43: 133–138. doi:10.1042/BST20140217

6. Song Y, Ke Y, Kang M, Bao R. Function, molecular mechanisms, and therapeutic potential of bacterial HtrA proteins: An evolving view. Comput Struct Biotechnol J. 2021;20: 40–49. doi:10.1016/j.csbj.2021.12.004

7. Kennedy M. Origin of PDZ (DHR, GLGF) domains. Trends in Biochemical Sciences. 1995;20: 350. doi:10.1016/S0968-0004(00)89074-X

8. Kim D-Y, Kim K-K. Structure and Function of HtrA Family Proteins, the Key Players in Protein Quality Control. BMB Reports. 2005;38: 266–274. doi:10.5483/BMBRep.2005.38.3.266

9. Xue R-Y, Liu C, Xiao Q-T, Sun S, Zou Q-M, Li H-B. HtrA family proteases of bacterial pathogens: pros and cons for their therapeutic use. Clinical Microbiology and Infection. 2021;27: 559–564. doi:10.1016/j.cmi.2020.12.017

10. Bongard J, Schmitz AL, Wolf A, Zischinsky G, Pieren M, Schellhorn B, et al. Chemical Validation of DegS As a Target for the Development of Antibiotics with a Novel Mode of Action. ChemMedChem. 2019;14: 1074–1078. doi:10.1002/cmdc.201900193

11. Krojer T, Sawa J, Schäfer E, Saibil HR, Ehrmann M, Clausen T. Structural basis for the regulated protease and chaperone function of DegP. Nature. 2008;453: 885–890. doi:10.1038/nature07004

12. Jiang J, Zhang X, Chen Y, Wu Y, Zhou ZH, Chang Z, et al. Activation of DegP chaperone-protease via formation of large cage-like oligomers upon binding to substrate proteins. Proc Natl Acad Sci USA. 2008;105: 11939–11944. doi:10.1073/pnas.0805464105

13. Ramsey DM, Wozniak DJ. Understanding the control of *Pseudomonas aeruginosa* alginate synthesis and the prospects for management of chronic infections in cystic fibrosis. Molecular Microbiology. 2005;56: 309–322. doi:10.1111/j.1365-2958.2005.04552.x

14. Hay ID, Schmidt O, Filitcheva J, Rehm BHA. Identification of a periplasmic AlgK– AlgX–MucD multiprotein complex in Pseudomonas aeruginosa involved in biosynthesis and regulation of alginate. Appl Microbiol Biotechnol. 2012;93: 215–227. doi:10.1007/s00253-011-3430-0

15. Wood LF, Ohman DE. Independent Regulation of MucD, an HtrA-Like Protease in Pseudomonas aeruginosa, and the Role of Its Proteolytic Motif in Alginate Gene Regulation. J Bacteriol. 2006;188: 3134–3137. doi:10.1128/JB.188.8.3134-3137.2006

16. Yorgey P, Rahme LG, Tan M, Ausubel FM. The roles of *mucD* and alginate in the virulence of *Pseudomonas aeruginosa* in plants, nematodes and mice. Molecular Microbiology. 2001;41: 1063–1076. doi:10.1046/j.1365-2958.2001.02580.x

17. Mochizuki Y, Suzuki T, Oka N, Zhang Y, Hayashi Y, Hayashi N, et al. *Pseudomonas aeruginosa* MucD Protease Mediates Keratitis by Inhibiting Neutrophil Recruitment and Promoting Bacterial Survival. Invest Ophthalmol Vis Sci. 2014;55: 240. doi:10.1167/iovs.13-13151

18. Damron FH, Yu HD. Pseudomonas aeruginosa MucD Regulates the Alginate Pathway through Activation of MucA Degradation via MucP Proteolytic Activity. J Bacteriol. 2011;193: 286–291. doi:10.1128/JB.01132-10

19. Gheorghita AA, Li YE, Kitova EN, Bui DT, Pfoh R, Low KE, et al. Structure of the AlgKX modification and secretion complex required for alginate production and biofilm attachment in Pseudomonas aeruginosa. Nat Commun. 2022;13: 7631. doi:10.1038/s41467-022-35131-6

20. Kim JH, Lee GH, Jeong J-H, Kim Y-G, Park HH. The structure of MucD from Pseudomonas syringae revealed N-terminal loop-mediated trimerization of HtrA-like serine protease. Biochemical and Biophysical Research Communications. 2023;688: 149175. doi:10.1016/j.bbrc.2023.149175

21. Keiski C-L, Harwich M, Jain S, Neculai AM, Yip P, Robinson H, et al. AlgK Is a TPR-Containing Protein and the Periplasmic Component of a Novel Exopolysaccharide Secretin. Structure. 2010;18: 265–273. doi:10.1016/j.str.2009.11.015

22. Riley LM, Weadge JT, Baker P, Robinson H, Codée JDC, Tipton PA, et al. Structural and Functional Characterization of Pseudomonas aeruginosa AlgX. J Biol Chem. 2013;288: 22299–22314. doi:10.1074/jbc.M113.484931

23. Bravo-Rodriguez K, Hagemeier B, Drescher L, Lorenz M, Rey J, Meltzer M, et al. Utilities for Mass Spectrometry Analysis of Proteins (UMSAP): Fast post-processing of mass spectrometry data. Rapid Comm Mass Spectrometry. 2018;32: 1659–1667. doi:10.1002/rcm.8243

24. Schechter I, Berger A. On the size of the active site in proteases. I. Papain. Biochemical and Biophysical Research Communications. 1967;27: 157–162. doi:10.1016/S0006-291X(67)80055-X

25. Schillinger J, Koci M, Bravo-Rodriguez K, Heilmann G, Kaschani F, Kaiser M, et al. High resolution analysis of proteolytic substrate processing. J Biol Chem. 2024;300: 107812. doi:10.1016/j.jbc.2024.107812

26. Hedstrom L. Serine Protease Mechanism and Specificity. Chem Rev. 2002;102: 4501– 4524. doi:10.1021/cr000033x

27. Harkness RW, Ripstein ZA, Di Trani JM, Kay LE. Flexible Client-Dependent Cages in the Assembly Landscape of the Periplasmic Protease-Chaperone DegP. J Am Chem Soc. 2023;145: 13015–13026. doi:10.1021/jacs.2c11849

28. Qiao Z, Yokoyama T, Yan X-F, Beh IT, Shi J, Basak S, et al. Cryo-EM structure of the entire FtsH-HflKC AAA protease complex. Cell Reports. 2022;39: 110890. doi:10.1016/j.celrep.2022.110890

29. Bai X, Pan X, Wang X, Ye Y, Chang L, Leng D, et al. Characterization of the Structure and Function of Escherichia coli DegQ as a Representative of the DegQ-like Proteases of Bacterial HtrA Family Proteins. Structure. 2011;19: 1328–1337. doi:10.1016/j.str.2011.06.013

30. Kim S, Sauer RT. Cage assembly of DegP protease is not required for substrate-dependent regulation of proteolytic activity or high-temperature cell survival. Proc Natl Acad Sci USA. 2012;109: 7263–7268. doi:10.1073/pnas.1204791109

31. Holyoake LV, Poole RK, Shepherd M. The CydDC Family of Transporters and Their Roles in Oxidase Assembly and Homeostasis. Advances in Microbial Physiology. Elsevier; 2015. pp. 1–53. doi:10.1016/bs.ampbs.2015.04.002

32. Knutson CA, Jeanes A. A new modification of the carbazole analysis: Application to heteropolysaccharides. Analytical Biochemistry. 1968;24: 470–481. doi:10.1016/0003-2697(68)90154-1

33. Wozniak DJ, Wyckoff TJO, Starkey M, Keyser R, Azadi P, O’Toole GA, et al. Alginate is not a significant component of the extracellular polysaccharide matrix of PA14 and PAO1 *Pseudomonas aeruginosa* biofilms. Proc Natl Acad Sci USA. 2003;100: 7907– 7912. doi:10.1073/pnas.1231792100

34. Shen M, Zhang H, Shen W, Zou Z, Lu S, Li G, et al. Pseudomonas aeruginosa MutL promotes large chromosomal deletions through non-homologous end joining to prevent bacteriophage predation. Nucleic Acids Research. 2018;46: 4505–4514. doi:10.1093/nar/gky160

35. Dar D, Sorek R. Extensive reshaping of bacterial operons by programmed mRNA decay. Buchrieser C, editor. PLoS Genet. 2018;14: e1007354. doi:10.1371/journal.pgen.1007354

36. Schurr MJ, Martin DW, Mudd MH, Hibler NS, Boucher JC, Deretic V. The algD promoter: regulation of alginate production by Pseudomonas aeruginosa in cystic fibrosis. Cell Mol Biol Res. 1993;39: 371–376.

37. Low KE, Gheorghita AA, Tammam SD, Whitfield GB, Li YE, Riley LM, et al. Pseudomonas aeruginosa AlgF is a protein–protein interaction mediator required for acetylation of the alginate exopolysaccharide. J Biol Chem. 2023;299: 105314. doi:10.1016/j.jbc.2023.105314

38. Baker P, Ricer T, Moynihan PJ, Kitova EN, Walvoort MTC, Little DJ, et al. P. aeruginosa SGNH Hydrolase-Like Proteins AlgJ and AlgX Have Similar Topology but Separate and Distinct Roles in Alginate Acetylation. Parsek MR, editor. PLoS Pathog. 2014;10: e1004334. doi:10.1371/journal.ppat.1004334

39. Krojer T, Garrido-Franco M, Huber R, Ehrmann M, Clausen T. Crystal structure of DegP (HtrA) reveals a new protease-chaperone machine. Nature. 2002;416: 455–459. doi:10.1038/416455a

40. Władyka B, Pustelny K. Regulation of bacterial protease activity. Cellular and Molecular Biology Letters. 2008;13. doi:10.2478/s11658-007-0048-4

41. Nagy ZA, Szakács D, Boros E, Héja D, Vígh E, Sándor N, et al. Ecotin, a microbial inhibitor of serine proteases, blocks multiple complement dependent and independent microbicidal activities of human serum. Mitchell TJ, editor. PLoS Pathog. 2019;15: e1008232. doi:10.1371/journal.ppat.1008232

42. Jones CH, Dexter P, Evans AK, Liu C, Hultgren SJ, Hruby DE. *Escherichia coli* DegP Protease Cleaves between Paired Hydrophobic Residues in a Natural Substrate: the PapA Pilin. J Bacteriol. 2002;184: 5762–5771. doi:10.1128/JB.184.20.5762-5771.2002

43. The UniProt Consortium, Bateman A, Martin M-J, Orchard S, Magrane M, Adesina A, et al. UniProt: the Universal Protein Knowledgebase in 2025. Nucleic Acids Research. 2025;53: D609–D617. doi:10.1093/nar/gkae1010

44. Punjani A, Rubinstein JL, Fleet DJ, Brubaker MA. cryoSPARC: algorithms for rapid unsupervised cryo-EM structure determination. Nat Methods. 2017;14: 290–296. doi:10.1038/nmeth.4169

45. Jumper J, Evans R, Pritzel A, Green T, Figurnov M, Ronneberger O, et al. Highly accurate protein structure prediction with AlphaFold. Nature. 2021;596: 583–589. doi:10.1038/s41586-021-03819-2

46. Emsley P, Lohkamp B, Scott WG, Cowtan K. Features and development of *Coot*. Acta Crystallogr D Biol Crystallogr. 2010;66: 486–501. doi:10.1107/S0907444910007493

47. Adams PD, Afonine PV, Bunkóczi G, Chen VB, Davis IW, Echols N, et al. *PHENIX*: a comprehensive Python-based system for macromolecular structure solution. Acta Crystallogr D Biol Crystallogr. 2010;66: 213–221. doi:10.1107/S0907444909052925

48. Goddard TD, Huang CC, Meng EC, Pettersen EF, Couch GS, Morris JH, et al. UCSF ChimeraX: Meeting modern challenges in visualization and analysis. Protein Science. 2018;27: 14–25. doi:10.1002/pro.3235

## Supplementary References

1. Tian Z, Cheng S, Xia B, Jin Y, Bai F, Cheng Z, et al. Pseudomonas aeruginosa ExsA Regulates a Metalloprotease, ImpA, That Inhibits Phagocytosis of Macrophages. Bäumler AJ, editor. Infect Immun. 2019;87: e00695–19. doi:10.1128/IAI.00695-19

2. Seviour T, Winnerdy FR, Wong LL, Shi X, Mugunthan S, Foo YH, et al. The biofilm matrix scaffold of Pseudomonas aeruginosa contains G-quadruplex extracellular DNA structures. npj Biofilms Microbiomes. 2021;7: 27. doi:10.1038/s41522-021-00197-5

3. Huang W, Wilks A. A rapid seamless method for gene knockout in Pseudomonas aeruginosa. BMC Microbiol. 2017;17: 199. doi:10.1186/s12866-017-1112-5

4. Lee MD, Henk AD. RSF1010-based shuttle vectors for cloning and expression in Pasteurella multocida. Veterinary Microbiology. 1997;54: 369–374. doi:10.1016/S0378-1135(96)01294-1

5. Yan X, Yang C, Wang M, Yong Y, Deng Y, Gao Y. Structural analyses of the AAA+ ATPase domain of the transcriptional regulator GtrR in the BDSF quorum-sensing system in *Burkholderia cenocepacia*. FEBS Letters. 2022;596: 71–80. doi:10.1002/1873-3468.14244

